# A continental-scale analysis reveals the latitudinal gradient of stomatal density across amphistomatous species: Evolutionary history vs. present-day environment

**DOI:** 10.1101/2023.10.21.563406

**Authors:** Congcong Liu, Kexiang Huang, Yifei Zhao, Ying Li, Nianpeng He

**Affiliations:** Key Laboratory of Ecology and Environment in Minority Areas (Minzu University of China), National Ethnic Affairs Commission, Beijing 100081, China; College of Life and Environmental Sciences, Minzu University of China, Beijing 100081, China; Key Laboratory of Ecosystem Network Observation and Modeling, Institute of Geographic Sciences and Natural Resources Research, Chinese Academy of Sciences, Beijing 100101, China; Center for Ecological Research, Northeast Forestry University, Harbin 150040, China; Earth Critical Zone and Flux Research Station of Xing’an Mountains, Chinese Academy of Sciences, Daxing’anling 165200, China

**Keywords:** stomatal density, stomatal ratio, growth form, amphistomy, adaptation, evolutionary history

## Abstract

Amphistomy is a potential method for increasing photosynthetic rate; however, latitudinal gradients of stomatal density across amphistomatous species and their drivers remain unknown. Here, the adaxial stomatal density (SD_ad_) and abaxial stomatal density (SD_ab_) of 486 amphistomatous species-site combinations, belonging to 32 plant families, were collected from China, and their total stomatal density (SD_total_) and stomatal ratio (SR) were calculated. Overall, these four stomatal traits did not show significant phylogenetic signals. There were no significant differences in SD_ab_ and SD_total_ between woody and herbaceous species, but SD_ad_ and SR were higher in woody species than in herbaceous species. Besides, a significantly positive relationship between SD_ab_ and SD_ad_ was observed. We also found that stomatal density (including SD_ab_, SD_ad_, and SD_total_) decreased with latitude while SR increased with latitude, and temperature seasonality was the most important environmental factor driving it. Besides, evolutionary history (represented by both phylogeny and species) explained about 10–22 fold more of the variation in stomatal traits than the present-day environment (65.2%–71.1% vs. 2.9%–6.8%). Our study extended our knowledge of trait-environment relationships and highlighted the importance of evolutionary history in driving stomatal trait variability.

## Introduction

Stomata are small pores on leaf surfaces and stalks, surrounded by a pair of guard cells. Their primary functions include regulating leaf gas exchange between the atmosphere and the chloroplast and optimizing carbon gain per unit water loss (M. Haworth, Marino, Loreto, & Centritto, 2021; Hetherington & Woodward, 2003; Raven, 2002). Stomatal control is achieved via dynamic opening and closing at a short time scale and morphology adjustment, including the density and size of stomata, at a long time scale (Franks & Farquhar, 2006). For most terrestrial species, stomata are located only on the lower leaf surface (hypostomy); however, the presence of stomata on both leaf sides (amphistomy) is not unusual since it is adaptively beneficial to plants, particularly herbaceous growth forms that are exposed to greater light intensities and lower precipitation circumstances (Mott, Gibson, & O’leary, 1982; Christopher D. Muir, 2015). Although amphistomy has garnered the attention of ecologists and botanists (Drake, de Boer, Schymanski, & Veneklaas, 2019; M. Haworth et al., 2021; Wall et al., 2022), variation in the stomatal density of amphistomatous species and their environmental drivers at large scales have not been investigated.

Amphistomatous species typically have higher gaseous exchange capacities than hypostomatous species (Xiong & Flexas, 2020), owing to stomata on the adaxial leaf surface that provide the benefits of shortening the transport distance of CO_2_ between leaf interiors and the atmosphere and reducing CO_2_ diffusion resistance by preventing condensation of water (Buckley, John, Scoffoni, & Sack, 2017; Drake et al., 2019). In addition, several plant families exhibit distinct evolutionary trends shifting from hypostomy to amphistomy. As a result, amphistomy is thought to be more evolved than hypostomy (Mott et al., 1982). Here are three possible scenarios that could illustrate how plant species transformed from hypostomaty into amphistomaty (Christopher D. Muir, 2018). In scenario 1, plants could hold their abaxial stomatal density (SD_ab_) constant, but if they add new stomata to their adaxial leaf surface, then their total stomatal density (SD_total_) would increase as well; under this scenario, SD_ab_ would not be correlated with adaxial stomatal density (SD_ad_), and the stomatal ratio (SR, the ratio of SD_ab_ to SD_ad_) would be mainly determined by SD_ad_. In scenario 2, plants could keep SD_total_ constant while moving abaxial stomata into their adaxial leaf surface; in this case, SD_ab_ would be negatively correlated with SD_ad_, and both SD_ab_ and SD_ad_ mediate SR change. In scenario 3, if both SD_ab_ and SD_ad_ increase, then SD_total_ would increase as well; under this scenario, SD_ab_ would be positively correlated with SD_ad_, and both SD_ab_ and SD_ad_ mediate change in SR. Theoretically, scenario 3 was the most effective way to improve the leaf gas exchange capacity. Besides, previous studies have reported that SD_ab_ and SD_ad_ are positively correlated using 29 amphistomatous species (Xiong & Flexas, 2020), which also suggested that Scenario 3 might be the strategy adopted by plants. However, it has not been rigorously tested on a broad phylogenetic scale.

The functional trait-environment relationship has been a central issue in ecological research for more than a century (Bruelheide et al., 2018). How climate and soil drive the variability of plant economic traits and size-related traits has been explored on a global scale (Boonman et al., 2020; Joswig et al., 2022). Now, a growing number of studies have focused on the relationship between stomatal traits and the environment, such as the fact that mean annual precipitation and temperature are the main drivers of stomatal density across 737 forest species (Liu et al., 2018); mean annual temperature and precipitation are the main drivers of stomatal density in *Pinus sylvestris* (Marek et al., 2022) and *Quercus variabilis* (Du, Zhu, Kang, & Liu, 2021). Based on the above studies, we inferred that mean annual temperature and precipitation might also be important factors in determining stomatal density across amphistomatous species. Although stomata on the upper leaf surface contribute a second parallel pathway for gaseous exchange (Christopher D. Muir, 2015), amphistomy might be costly (Christopher D. Muir, 2018; Christopher D Muir, 2019). Stomata on the adaxial leaf surface increased risk of infection (McKown et al., 2014). In dry and high solar intensity conditions, fungi were less able to survive and reproduce; therefore, plants could develop more stomata on the adaxial surface to maximize their productivity without the threat of fungi. Therefore, we hypothesized that drought and high radiation intensity would support species with higher SD_ad_and/or SR.

In this study, we aimed to collect the stomatal density of amphistomatous species from China to answer the following questions: (1) What is the relationship between SD_ab_ and SD_ad_, and how do SD_ab_ and SD_ad_ determine the change in SR? (2) Does the stomatal density of amphistomatous species show latitudinal patterns, and what are the driving factors? We inferred that such latitudinal patterns were jointly regulated by both current environmental factors (climate and soil variables) and evolutionary history, and the evolutionary history might explain more variability in the stomatal density of amphistomatous species than current environmental factors, due to increasing studies that have found plant trait variations are better explained by evolutionary history than by environmental factors (Sardans et al., 2021; Yan et al., 2023). China has almost all kinds of vegetation found in the Northern Hemisphere, including cold-temperate coniferous forest, temperate coniferous and broad-leaved mixed forest, warm temperate deciduous broad-leaved forest, subtropical evergreen broad-leaved forest, tropical rain forest, temperate steppe, temperate desert, and alpine vegetation of the Qinghai-Tibetan Plateau. Furthermore, Chinese ecologists and botanists have long been interested in stomatal traits, and there have been numerous studies on the topic published in either Chinese or English. Due to these exogenous factors, it was possible to integrate the fragmented data and create a database that detailed the stomatal density of amphistomatous species along climatic gradients. Finally, the stomatal densities of 486 amphistomatous species-site combinations, which were evenly distributed throughout all vegetation types, were collected to address the aforementioned research question.

## Materials and Methods

### Literature survey, stomatal traits, and environmental factors

To examine the drivers of stomatal density across diverse amphistomatous species at a regional scale, we first compiled abaxial stomatal density (SD_ab_) and adaxial stomatal density (SD_ad_) data from peer-reviewed literature and dissertations of Chinese students. The keywords “stomata traits,” “stomatal density,” and “stomatal ratio” were used to retrieve potential literature from the Web of Science (www.webofknowledge.com) and the China National Knowledge Infrastructure database (http://epub.cnki.net). Publications that were included in our meta-analysis met the following criteria: 1) The name of the amphistomatous species must be known, and SD_ab_ and SD_ad_ were reported; 2) Plant samples had to be collected in the wild, with no experimental treatments or indoor cultures; 3) Stomatal density was measured from healthy mature plant leaves; 4) The number of the stomata was manually counted. Moreover, stomatal trait values for each site-species combination were recorded when data for one species were collected from multiple sites. In most cases, stomatal density data was available in the table. When stomatal traits were presented only in figures, we used the WebPlotDigitizer software (https://automeris.io/WebPlotDigitizer/) to extract them. The geographical coordinates of the sampling sites were also recorded to obtain the corresponding climate and soil variables. The preferred reporting items for systematic reviews and meta-analyses (PRISMA) flowchart of the publication selection procedure is shown in Fig. S1. The final dataset included 486 amphistomatous species-site combinations, belonging to 32 families, 86 genera and 291 species (Fig. 1). Specifically, 418 species-site combinations have coordinates and 68 species-site combinations do not have exact coordinates (Table S1). Total stomatal density (SD_total_) and stomatal ratio (SR) were calculated as:

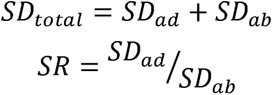

**Fig.1.**
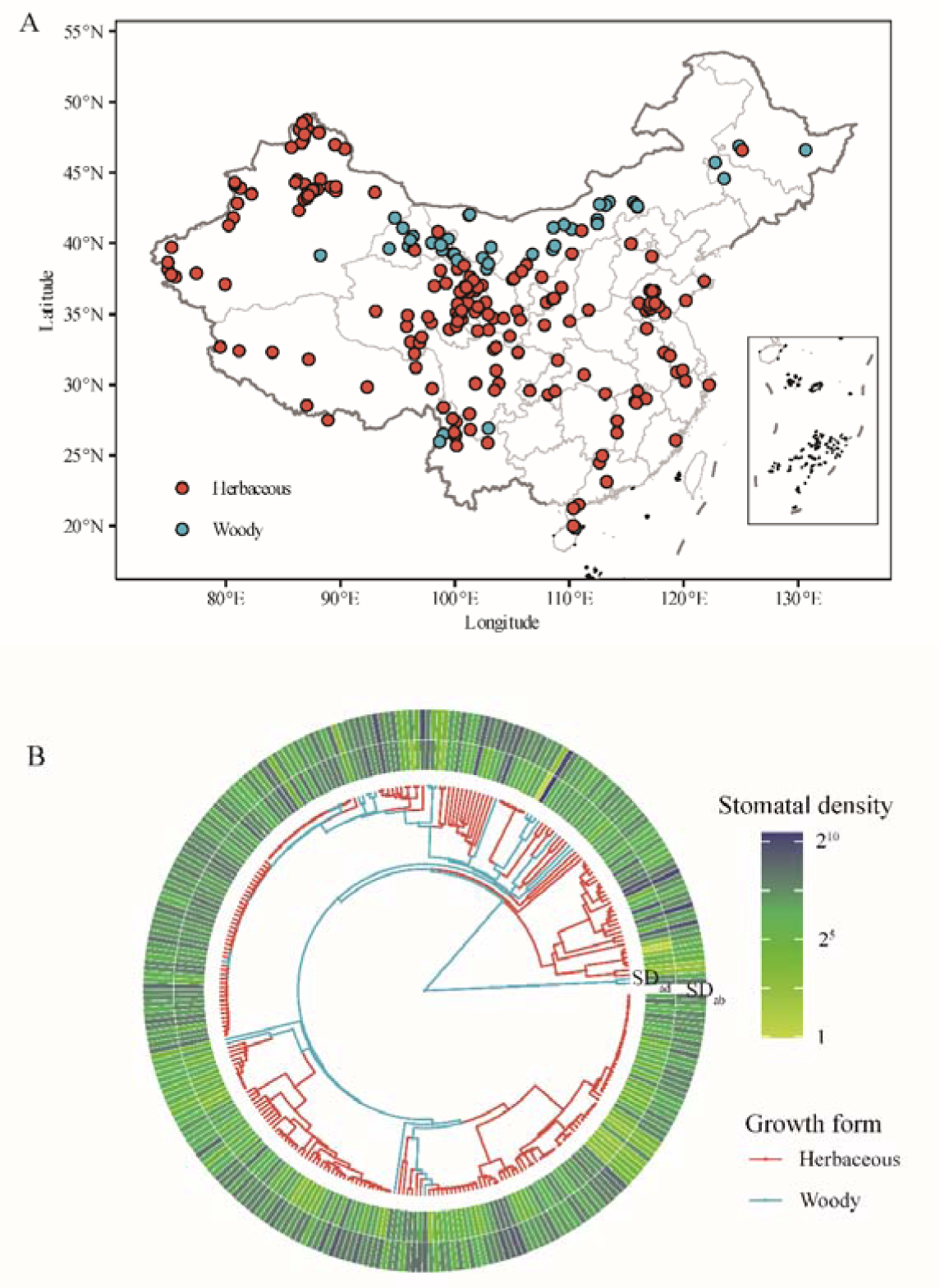
The distribution of the 242 sampling sites (A) and stomatal density of amphistomatous species along plant phylogeny (B). Provincial boundaries are shown on the map of China. There are 291 species, 86 genera, and 32 families represented in the phylogenetic tree. SD_ad_, adaxial stomatal density; SD_ab_, abaxial stomatal density.

All environmental variables were extracted from databases using the function ‘extract’ in the R package raster. Mean annual temperature (MAT), temperature seasonality (TS), mean annual precipitation (MAP), precipitation seasonality (PS), and solar radiation were obtained from WorldClim version 2.1 (http://www.worldclim.org/). The aridity index (AI, the ratio of mean annual precipitation to potential evapotranspiration) was obtained from the Consortium of Spatial Information (CGIAR-CSI) website (https://cgiarcsi.community/). All climate variables were based on average values from 1950 to 2000 using weather station data at 0.5° × 0.5° resolution. Properties of topsoil (e.g., nitrogen content, pH, bulk density, and soil texture) were collected from the SoilGrids (https://soilgrids.org/). Overall, the six climatic variables and five soil variables used in this study are provided in Table S2.

### Phylogeny construction and phylogenetic signal

The scientific names of plant species were corrected using the “Taxonstand” package of R (Cayuela, Granzow-de la Cerda, Albuquerque, & Golicher, 2012), which used the online search engine of The Plant List (http://www.theplantlist.org) to retrieve information on the latest taxonomic status of each species. Phylogenies of plant species were generated using the “V.PhyloMaker” package of R (Jin & Qian, 2019), which includes the largest dated phylogeny of vascular plants. The final tree was ultrametric with time-calibrated branches. Note that we constructed two phylogenetic trees: one included 291 plant species (the result of removing all duplicates from 486 records), and the other included 257 plant species (the result of removing all duplicates from 418 records). The former was used to calculate phylogenetic signals, test the differences between plant growth forms, and explore the relationship between adaxial and abaxial stomatal density; the latter was used to test the relationships of stomatal traits to latitude and environmental factors.

Before calculating phylogenetic signals, stomatal traits were averaged for each plant species. We used Blomberg’s K (Blomberg, Garland JR., & Ives, 2003) to evaluate phylogenetic signals using species-mean values. Although polytomies have negligible effect on these indices, we used the *multi2di* function of the “ape” package of R (Paradis & Schliep, 2018) to transform polytomies into dichotomous trees before estimating phylogenetic signals (Ibanez et al., 2021; Münkemüller et al., 2012). Blomberg’s K was calculated using the *Phylosig* function of the “phytools” package (Revell, 2012). K = 0 suggested absence of a phylogenetic signal, K = 1 indicated that the trait variation conformed to the Brownian model, and K > 1 indicated very strong phylogenetic conservatism.

### Graphical and statistical analyses

SD_ab_ and SD_ad_ matched with phylogenetic trees were generated using the “ggtree” package (Yu, Smith, Zhu, Guan, & Lam, 2017). Phylogenetic relatedness was a significant source of non-independence between species, and some species appeared at more than one site in this dataset; therefore, all the analyses were conducted using Bayesian phylogenetic linear mixed models with phylogeny and species as random factors. Bayesian phylogenetic linear mixed models were performed using the R package *MCMCglmm* (Hadfield, 2010).

All species were classified into woody and herbaceous species based on their descriptions in the original publication, Flora of China (http://www.iplant.cn/). To compare stomatal traits between growth forms (woody vs. herbaceous), “growth form” was taken as a fixed factor in the Bayesian phylogenetic linear mixed models, and the code was written as: MCMCglmm.updateable(Traits∼growth, random=∼ phylogeny + species). If 95% credible intervals of regression coefficient excluded zero, indicating that the stomatal trait was significantly different between woody and herbaceous species. To explore the relationship between SD_ab_ and SD_ad_, the code was written as: MCMCglmm.updateable(SD_ad_ ∼ SD_ab_, random=∼ phylogeny + species). To explore how SD_ab_ and SD_ad_ determined SR, SD_ab_ and SD_ad_ were first standardized (mean =0, standard deviation =1), then the code was written as: MCMCglmm.updateable(SR ∼ SD_ad_ + SD_ab_, random=∼ phylogeny + species). If the absolute values of the regression coefficients of SD_ad_ and SD_ab_ did not show a significant difference, it indicated that SD_ad_ and SD_ab_ played an equally important role in determining SR. To explore the relationships of stomatal traits to latitude and environmental factors, the code was written as: MCMCglmm.updateable (Trait ∼latitude or environmental factor, random=∼ phylogeny + species). Bayesian phylogenetic linear mixed models were also used to disentangle the relative contributions of current environmental factors and evolutionary history to stomatal trait variability, and the code was written as: MCMCglmm.updateable (Trait∼ environmental factor1+ environmental factor2+…, random=∼ phylogeny + species).

All graphical and statistical analyses were performed using R software (http://www.R-project.org), and the data and R code were provided in Supplementary Information.

## Results

### Phylogenetic signals of stomatal traits

Blomberg’s K was used to evaluate the phylogenetic signals of stomatal traits across 291 amphistomatous species. Overall, Blomberg’s K values of SD_ab_, SD_ad_, SD_total_, and SR were very small (ranging from 0.001 to 0.007) and did not significantly differ from 0 (Table 1).

**Table 1.** Phylogenetic conservatism indices across 291 amphistomatous species.

| Stomatal traits | Blomberg's K | P value |
| --- | --- | --- |
| Adaxial stomatal density ($SD_{ad}$) | 0.007 | 0.380 |
| Abaxial stomatal density ($SD_{ab}$) | 0.007 | 0.404 |
| Total stomatal density ($SD_{total}$) | 0.006 | 0.514 |
| Stomatal ratio (SR) | 0.001 | 0.901 |

### Stomatal trait variations and their differences between growth forms

Both SD_ab_ and SD_ad_ varied markedly across 486 amphistomatous species-site combinations (Fig. 2). We observed > 1440- and 223-fold variations in SD_ad_ and SD_ab_, respectively. Furthermore, SD_ad_ (135.82 ± 8.52 stomata mm^-2^) was lower than SD_ab_ (174.59 ± 8.58 stomata mm^-2^), varying from 0.967 to 1396 stomata mm^-2^ and from 4.62 to 1033 stomata mm^-2^, respectively (Table S3). The Bayesian phylogenetic linear mixed models showed that SD_ab_ and SD_total_ were not significantly different between woody and herbaceous species, while SD_ad_ and SR of woody species were much higher than those of herbaceous species (Table S4-S7).

**Fig.2.**
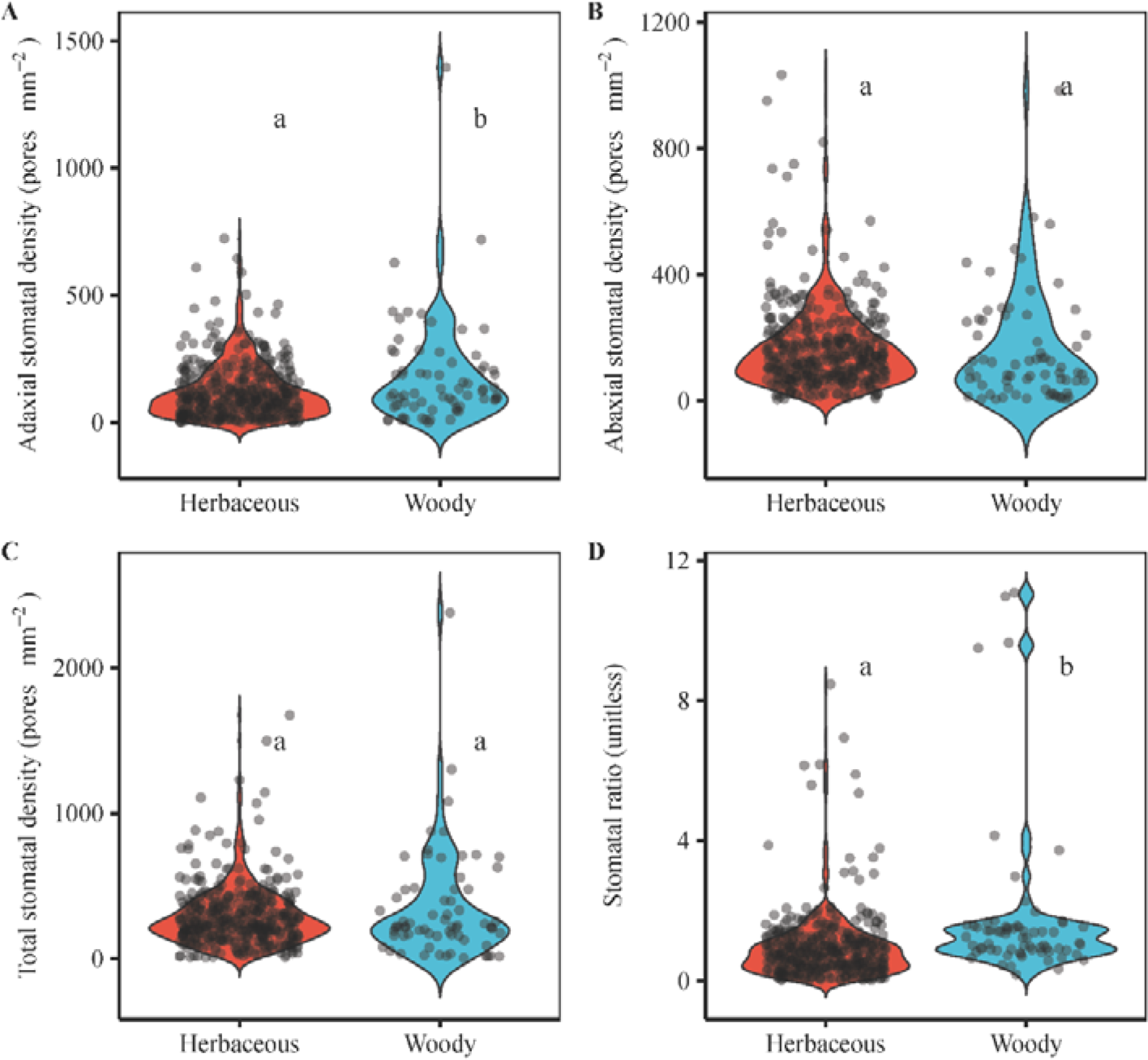
Distributions of stomatal trait for woody and herbaceous plants across amphistomatous species. Each point represents one species-site combination. The background violin plot characterizes the distribution of points in each plant growth form. All the statistics are estimated using the Bayesian phylogenetic linear mixed model with phylogeny and species as random factors. Same lowercase letters denote no significant difference, and different lowercase letters denote significant difference (*p* < 0.05). More details are provided in Table S4-S7.

### Relationships between adaxial and abaxial stomatal density

SD_ab_ and SD_ad_ were positively correlated with each other (R^2^m = 0.242, R^2^c = 0.875; Fig. 3 and Table S8). Absolute values of the regression coefficients of SD_ad_ and SD_ab_ in the multiple regression (SR∼ SD_ad_ + SD_ab_) were 0.57 and 0.61, respectively (Table S9), and their 95% confidence intervals were overlapped, indicating that SD_ad_ and SD_ab_ played an equally important role in determining SR.

**Fig.3.**
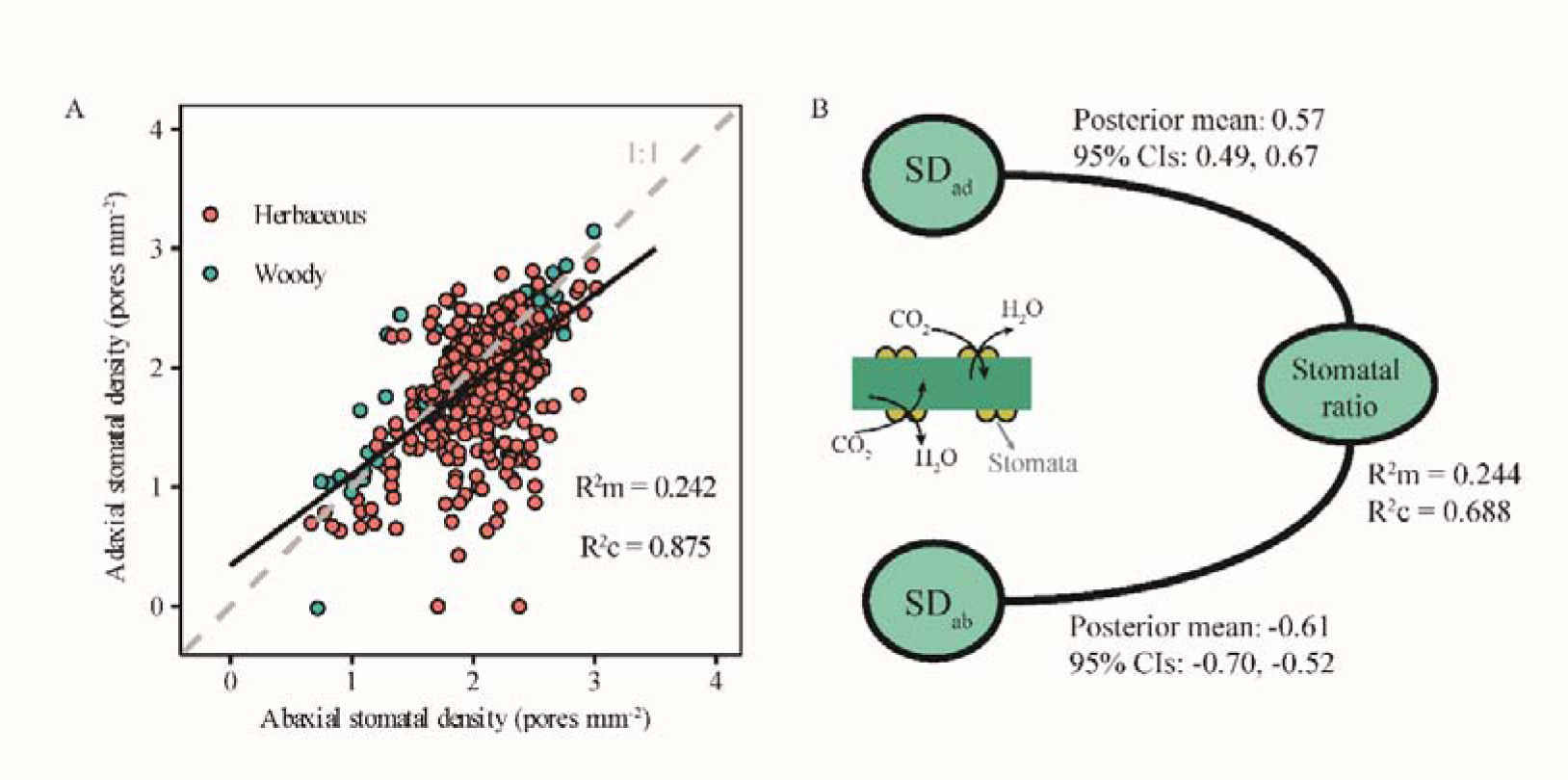
Relationships between abaxial and adaxial stomatal density (A) and their determinations on stomatal ratio (B) across amphistomatous species. Stomatal traits were log_10_-transformed and standardized (mean =0, standard deviation =1) in Panel A and Panel B, respectively. All the statistics are estimated using the Bayesian phylogenetic linear mixed model with phylogeny and species as random factors. SD_ad_, adaxial stomatal density, SD_ab_ adaxial stomatal density. R^2^m, marginal R^2^ (fixed effects only); R^2^c, conditional R^2^ (both fixed and random effects). More details are provided in Table S8 and S9.

### Bivariate relationships of stomatal traits to latitude and environmental factors

SD_ab_, SD_ad_, and SD_total_ decreased with latitude, while SR increased with latitude (Fig.4 and Table S10-S13). Among 11 environmental factors, SD_ad_ and SR were only significantly correlated with temperature seasonality (Fig.5); SD_ab_ and SD_total_ were significantly correlated with the aridity index, mean annual temperature, mean annual precipitation, and temperature seasonality. Compared with the relationships of SD_ab_ and SD_total_ to aridity index, and mean annual temperature and precipitation, the relationships of SD_ab_ and SD_total_ to temperature seasonality were strongest. Overall, SD_ab_, SD_ad_, SD_total_, and SR were less influenced by soil properties (Table S14-S17).

**Fig.4.**
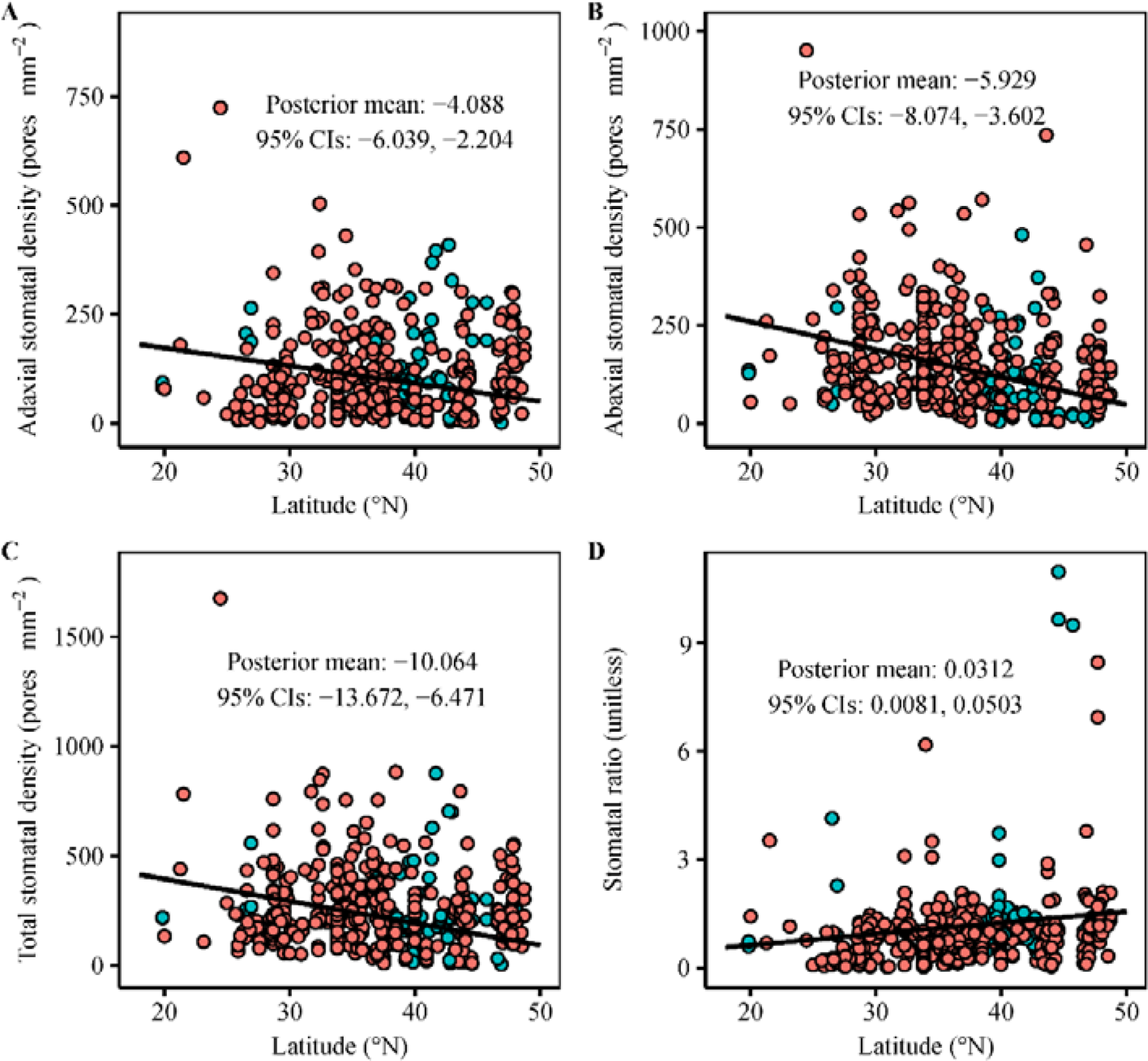
Latitudinal gradients of stomatal density across amphistomatous species. Pink point, herbaceous species; green point, woody species. The fitted lines are estimated using the Bayesian phylogenetic linear mixed model with phylogeny and species as random factors. More details were shown in Table S10-S13.

**Fig.5.**
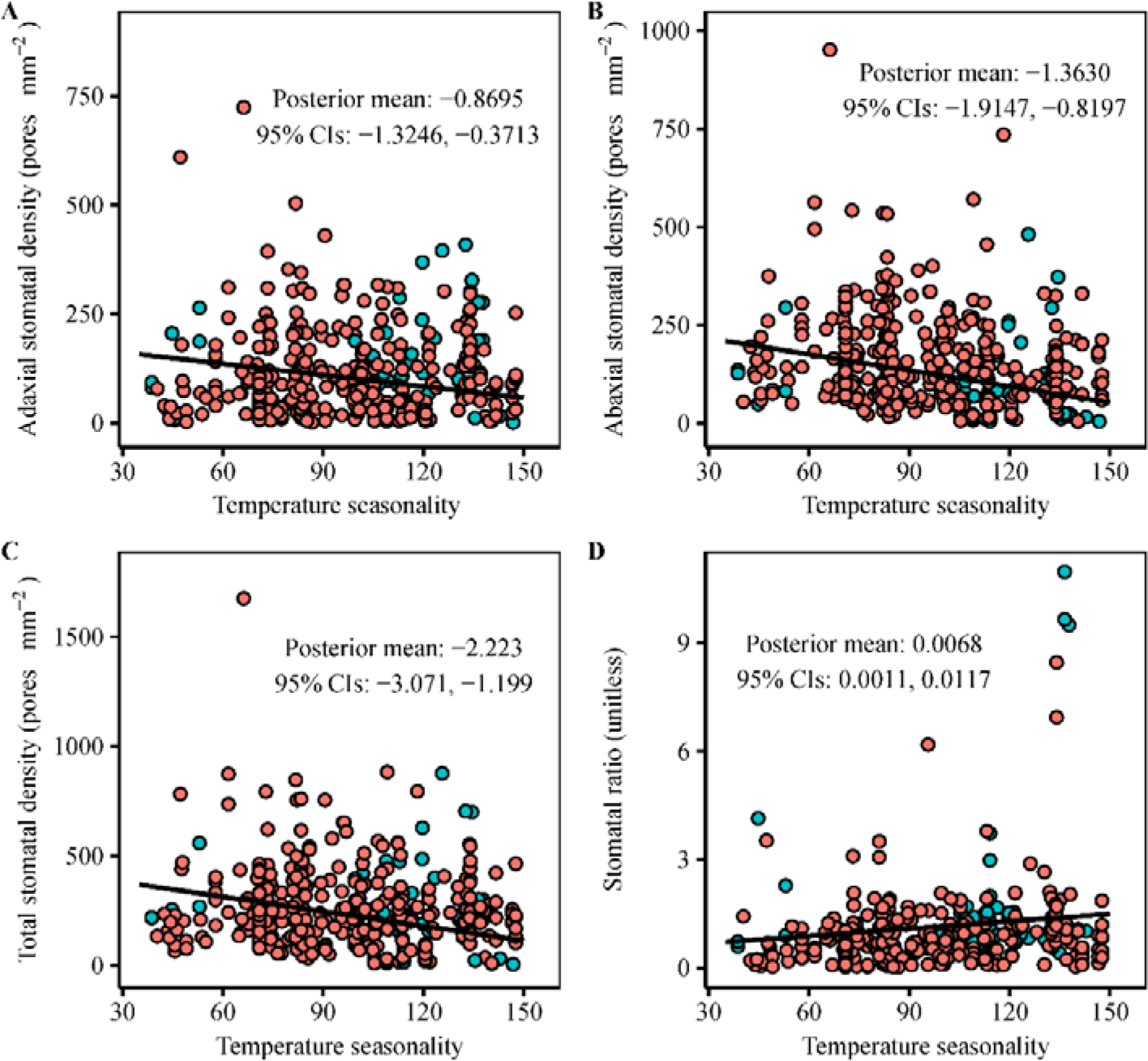
Relationships between stomatal traits and temperature seasonality. Pink point, herbaceous species; green point, woody species. The fitted lines are estimated using the Bayesian phylogenetic linear mixed model with phylogeny and species as random factors. More details were shown in Table S14-17.

### Relative contributions of environmental factors and evolutionary history to stomatal trait variability

Although SD_ab_, SD_ad_, SD_total_, and SR were significantly influenced by environmental factors, the explanations of environmental factors on stomatal traits were very low (ranging from 2.9% to 6.8%; Fig.6 and Table S18-S21). Phylogeny (accounted for variability in the shared ancestry) was the most important factor that determined stomatal traits and explained more than half of the variances of stomatal traits (ranging from 52.9% to 65.2%). Variances of stomatal traits explained by species (species-specific variance independent of the shared ancestry) were also not very high (ranging from 1.4% to 9.8%).

**Fig.6.**
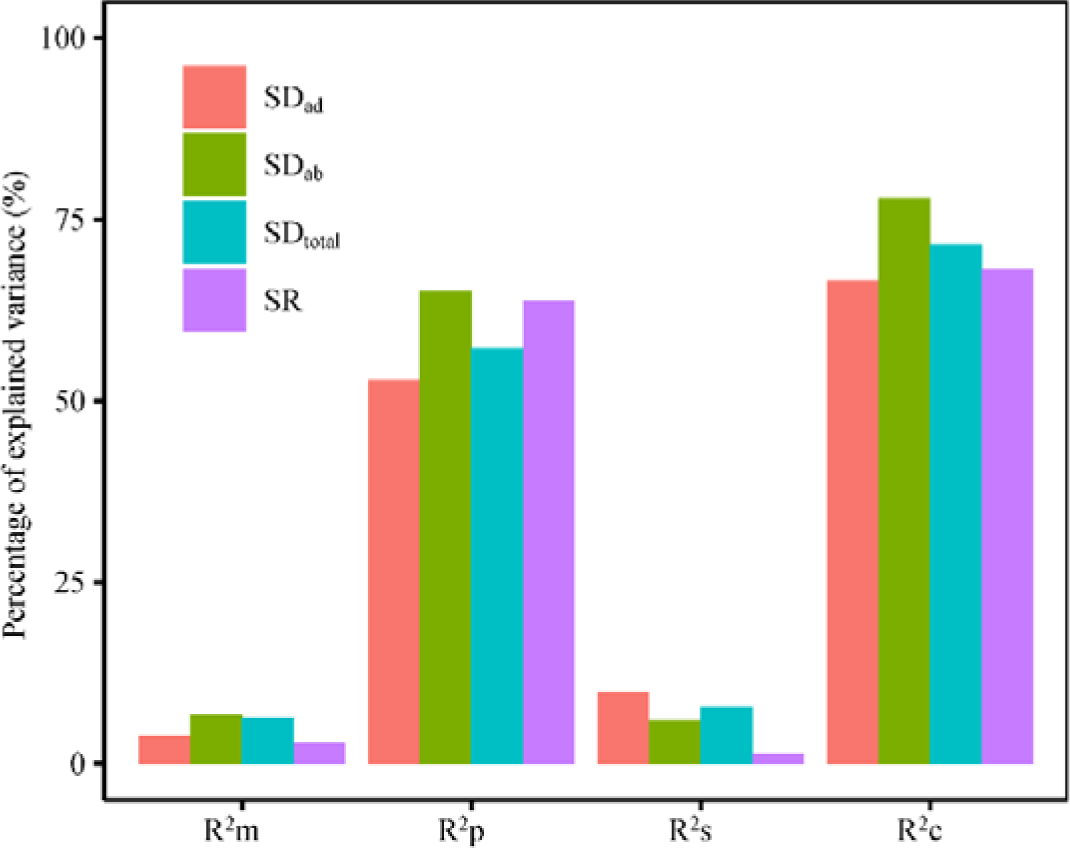
Variances of stomatal traits explained by environmental factors and evolutionary history (represented by both phylogeny and species). SD_ad_, adaxial stomatal density; SD_ab_, abaxial stomatal density; SD_total_, total stomatal density; SR, stomatal ratio. R^2^c, percentage of variance explained by both fixed (environmental factors) and random factors (evolutionary history); R^2^m, percentage of variance explained by the environmental factors; R^2^p, percentage of variance explained by phylogeny; R^2^s, percentage of variance explained by species. All the statistics are estimated using the Bayesian phylogenetic linear mixed model with phylogeny and species as random factors. More details were shown in Table S18-S21.

## Discussion

Changes in the occurrence of stomatal pores from one leaf side to both sides shortened the pathway for CO_2_ transport between leaf anatomy and the atmosphere (Parkhurst, 1978). Thus, amphistomy emerged as an important means to improve the photosynthetic rate (Matthew Haworth et al., 2018; Christopher D. Muir, 2018; Xiong & Flexas, 2020). To the best of our knowledge, this is the first study that explores the latitudinal patterns of stomatal traits (including SD_ab_, SD_ad_, SD_total_, and SR) and their environmental drivers across amphistomatous species at broad phylogenetic scales. Our dataset included stomatal density of 486 amphistomatous species-site combinations, which was larger than the stomatal dataset of Christopher D. Muir (2015) at the global scale. Fast-growing herbaceous annuals have been considered the most common amphistomatous species (Drake et al., 2019), but in our dataset, 222 out of 291 species were herbaceous perennials, indicating that not herbaceous annuals, but herbaceous perennials are the most common amphistomatous species in the plant kingdom. We also found stomatal density of amphistomatous species exhibited very weak or no phylogenetic signals (Table 1), indicating that the evolution of amphistomy was more flexible and less constrained by leaf epidermal space (de Boer et al., 2016).

### Positive relationship between SD_ab_ and SD_ad_ and their determinations in SR

We observed a positive correlation between SD_ab_ and SD_ad_ across all species, and such relationship was also observed within species (Fanourakis et al., 2014). These findings suggest that co-variation between SD_ab_ and SD_ad_ was ubiquitous, and genetic constraints, which resulted in evolutionary integration (Caetano & Harmon, 2017; Goswami, Binder, Meachen, & O’Keefe, 2015; Melo, Porto, Cheverud, & Marroig, 2016), might account for it. In other words, adjusting SD_total_ usually will be accompanied by simultaneous changes in SD_ab_ and SD_ad_. From an ecological perspective, independently varying traits would produce more putative plant trait combinations, thereby allowing plants more freedom to adjust to complex environments (Flores-Moreno et al., 2019). For example, leaf economic traits were decoupled with leaf hydraulic traits (Li et al., 2015) and stem economic traits (Baraloto et al., 2010), and these multiple trait combinations might more effectively allow species to adapt to various niche dimensions. Thus, compared with co-variation between SD_ab_ and SD_ad_, independent variability of SD_ab_ and SD_ad_ was more adaptive and cost-effective for plants. Previous studies found that stomata on upper and lower leaf surfaces can independently regulate gaseous exchange under fluctuating environmental conditions (Heichel & Anagnostakis, 1978; Pearson, Davies, & Mansfield, 1995; Richardson, Brodribb, & Jordan, 2017; Wall et al., 2022), which might compensate for the disadvantages of co-variation between SD_ab_ and SD_ad_.

The SR of leaf fossil records could be used to discriminate between open vegetation and closed forest, thus interpreting paleoenvironments (Jordan, Carpenter, & Brodribb, 2014). Furthermore, SR was also a guide for future directional breeding during domestication for increasing productivity and/or resource-use efficiency (Fanourakis et al., 2014; Milla, de Diego-Vico, & Martín-Robles, 2013). In the present study, we found that SD_ab_ and SD_ad_ played equal roles in mediating the change in SR. This conclusion was at odds with Christopher D. Muir (2018), possibly because this study only concentrated on amphistomatous species while the analysis of Muir (2018) included both amphistomatous and hypostomatous species. Amphistomy could improve photosynthesis even under similar SD_total_ (M. Haworth et al., 2021). Therefore, understanding how SD_ab_ and SD_ad_ determined SR was essential for crop improvement (Milla et al., 2013; Richardson et al., 2017).

### Latitudinal patterns of stomatal traits and their climatic drivers

Latitudinal patterns of stomatal density (including SD_ab_, SD_ad_, and SD_total_) were apparently observed, and stomatal density increased with latitude, and temperature seasonality, rather than mean annual precipitation and temperature, was the most important climatic driver for them (Fig. 5). Specifically, stomatal density was lower in regions with a higher temperature seasonality, and such results were very striking. The construction and maintenance of the stomata were expensive (de Boer et al., 2016); thus, plants need to weigh the cost and benefit of stomata under specific conditions. In areas with a small temperature seasonality, photosynthetic rates were always maintained in their optimal state, so plants could allocate more energy and matter to develop more stomata on the leaf surface, and higher stomatal density would also in turn boost photosynthesis. In areas with a large temperature seasonality, photosynthetic rates might decrease because of too high or too low temperatures, resulting in plants not developing enough stomata.

Contrary to our hypothesis that aridity index and solar intensity were the main drivers of SR, we did not observe significant relationships between SR, aridity index, and solar intensity. SR was the ratio of SD_ad_ to SD_ab_, and SD_ad_ and SD_ab_ were mainly determined by temperature seasonality; therefore, temperature seasonality was also the main driver of SR. SD_ad_ and SD_ab_ were positively correlated, and both of them decreased with latitude and temperature seasonality. However, SD_ab_ was more sensitive to latitude and temperature seasonality than SD_ad_, resulting in SR increasing with latitude and temperature seasonality.

Stomatal density is sensitive to changing CO_2_ concentrations, and an inverse correlation between stomatal density and CO_2_ concentration has been well-established. Thus, the stomatal density of fossil leaves has been widely used to infer palaeo-CO_2_ concentrations (Matthew Haworth, Elliott-Kingston, & McElwain, 2011; M. Haworth et al., 2021; Royer, 2001). In this study, clear climatic trends of stomatal traits across amphistomatous species were revealed; thus, these findings might also provide empirical evidence to infer paleo-climate transitions (especially temperature seasonality) in the future.

### Evolutionary history should be emphasized in shaping the stomatal trait variability

Understanding what drives such a large variability of stomatal traits at the regional scale is the key to exploring the mechanism of plant adaptation across amphistomatous species. Temperature seasonality was the main driver of stomatal traits, which indeed could help us understand their latitudinal patterns; e.g., species with high stomatal density inhabit tropical regions, corresponding to low temperature seasonality. However, when we considered all current environmental factors (including climate and soil), it only explained 2.9%–6.8% of stomatal trait variability, which was much lower than we expected. Our Bayesian phylogenetic linear mixed models showed that phylogeny and species (the combination of the two is called evolutionary history) explained about 10–22 fold more of the variation in stomatal traits than in the present-day environment (65.2%–71.1% vs. 2.9%–6.8%), highlighting the importance of evolutionary history in driving stomatal trait variability at a large scale. Previous studies have demonstrated that the variability of leaf element content (Sardans et al., 2021) and photosynthetic capacity (Yan et al., 2023) were also mainly explained by evolutionary history. All of them emphasized the importance of evolutionary history in predicting the variability of plant functional traits, and next-generation dynamic global vegetation models should incorporate evolutionary history to improve their simulations.

### Limitations of the study

In this study, we constructed a database of amphistomatous species’ stomatal density at a regional scale to investigate the relationships between stomatal density and environmental variables. Our data were collected from published literature, and even though we set strict screening criteria, there were still some uncertainties. First, stomatal density measurements were taken in different years and seasons, so fluctuations in local climate had an effect on stomatal density; second, different researchers had different artificial errors in sample collection and stomatal trait measurement; and third, the influence of local factors (including microclimate, fine-scale soil properties, topography, and even biotic factors) on stomatal traits was not taken into account. Given that interspecific trait variation was much greater than intraspecific trait variation and that species turnover played important roles in large-scale patterns of plant traits, our results were robust enough on the whole. With the development of high-throughput techniques and deep learning-based methods (Liang et al., 2022; Sakoda et al., 2019), future studies could overcome the above defects to explore the stomatal density-environment relationship across amphistomatous species on a broader, even global scale.

## Conclusions

By analyzing stomatal traits of 486 amphistomatous species-site combinations at a regional scale, we found that stomatal traits (including SD_ab_, SD_ad_, SD_total_, and SR) of amphistomatous species did not show significant phylogenetic signals. Stomatal traits of amphistomatous species showed apparent latitudinal patterns, and temperature seasonality was the most important climatic driver. Evolutionary history (represented by both phylogeny and species) explained 65.2%–71.1% of stomatal trait variability, while the present-day environment only explained 2.9%–6.8% of stomatal trait variability. This study improved our understanding of the adaptations of amphistomatous species at a regional scale and highlighted that evolutionary history should be emphasized in shaping plant functional trait variability.

## Author contributions

C.L. planned and designed the research; C.L., K.H., Y.Z. and Y.L. collected data, analyzed data and wrote the manuscript; N.H. revised the manuscript.

## Data availability

The data and R code that support the findings of this study are provided in the Supplementary Information.

## References

Baraloto, C., Timothy Paine, C. E., Poorter, L., Beauchene, J., Bonal, D., Domenach, A.-M., . . . Chave, J. (2010). Decoupled leaf and stem economics in rain forest trees. Ecology Letters, 13(11), 1338–1347. 10.1111/j.1461-0248.2010.01517.x

Blomberg, S. P., Garland JR., T., & Ives, A. R. (2003). Testing for phylogenetic signal in comparative data: Behavioral traits are more labile. Evolution, 57(4), 717–745. doi:10.1111/j.0014-3820.2003.tb00285.x

Boonman, C. C. F., Benítez-López, A., Schipper, A. M., Thuiller, W., Anand, M., Cerabolini, B. E. L., . . . Santini, L. (2020). Assessing the reliability of predicted plant trait distributions at the global scale. Global Ecology and Biogeography, 29(6), 1034–1051. 10.1111/geb.13086

Bruelheide, H., Dengler, J., Purschke, O., Lenoir, J., Jiménez-Alfaro, B., Hennekens, S. M., . . . Jandt, U. (2018). Global trait–environment relationships of plant communities. Nature Ecology & Evolution, 2(12), 1906–1917. doi:10.1038/s41559-018-0699-8

Buckley, T. N., John, G. P., Scoffoni, C., & Sack, L. (2017). The sites of evaporation within leaves. Plant Physiology, 173(3), 1763–1782. doi:10.1104/pp.16.01605 %J Plant Physiology

Caetano, D. S., & Harmon, L. J. (2017). ratematrix: An R package for studying evolutionary integration among several traits on phylogenetic trees. Methods in Ecology and Evolution, 8(12), 1920–1927. 10.1111/2041-210X.12826

Cayuela, L., Granzow-de la Cerda, Í., Albuquerque, F. S., & Golicher, D. J. (2012). taxonstand: An r package for species names standardisation in vegetation databases. Methods in Ecology and Evolution, 3(6), 1078–1083. 10.1111/j.2041-210X.2012.00232.x

de Boer, H. J., Price, C. A., Wagner-Cremer, F., Dekker, S. C., Franks, P. J., & Veneklaas, E. J. (2016). Optimal allocation of leaf epidermal area for gas exchange. New Phytologist, 210(4), 1219–1228. 10.1111/nph.13929

Drake, P. L., de Boer, H. J., Schymanski, S. J., & Veneklaas, E. J. (2019). Two sides to every leaf: water and CO_2_ transport in hypostomatous and amphistomatous leaves. New Phytologist, 222(3), 1179–1187. doi:10.1111/nph.15652

Du, B., Zhu, Y., Kang, H., & Liu, C. (2021). Spatial variations in stomatal traits and their coordination with leaf traits in Quercus variabilis across Eastern Asia. Science of The Total Environment, 789, 147757. 10.1016/j.scitotenv.2021.147757

Fanourakis, D., Giday, H., Milla, R., Pieruschka, R., Kjaer, K. H., Bolger, M., . . . Ottosen, C.-O. (2014). Pore size regulates operating stomatal conductance, while stomatal densities drive the partitioning of conductance between leaf sides. Annals of Botany, 115(4), 555–565. doi:10.1093/aob/mcu247

Flores-Moreno, H., Fazayeli, F., Banerjee, A., Datta, A., Kattge, J., Butler, E. E., . . . Reich, P. B. (2019). Robustness of trait connections across environmental gradients and growth forms. Global Ecology and Biogeography, 28(12), 1806–1826. 10.1111/geb.12996

Franks, P. J., & Farquhar, G. D. (2006). The mechanical diversity of stomata and its significance in gas-exchange control. Plant Physiology, 143(1), 78–87. doi:10.1104/pp.106.089367

Goswami, A., Binder, W. J., Meachen, J., & O’Keefe, F. R. (2015). The fossil record of phenotypic integration and modularity: A deep-time perspective on developmental and evolutionary dynamics. Proceedings of the National Academy of Sciences, 112(16), 4891–4896. doi:10.1073/pnas.1403667112

Hadfield, J. D. (2010). MCMC Methods for Multi-Response Generalized Linear Mixed Models: The MCMCglmm R Package. Journal of Statistical Software, 33(2), 1–22. doi:10.18637/jss.v033.i02

Haworth, M., Elliott-Kingston, C., & McElwain, J. C. (2011). Stomatal control as a driver of plant evolution. Journal of Experimental Botany, 62(8), 2419–2423. doi:10.1093/jxb/err086

Haworth, M., Marino, G., Loreto, F., & Centritto, M. (2021). Integrating stomatal physiology and morphology: evolution of stomatal control and development of future crops. Oecologia, 197(4), 867–883. doi:10.1007/s00442-021-04857-3

Haworth, M., Scutt, C. P., Douthe, C., Marino, G., Gomes, M. T. G., Loreto, F., . . . Centritto, M. (2018). Allocation of the epidermis to stomata relates to stomatal physiological control: Stomatal factors involved in the evolutionary diversification of the angiosperms and development of amphistomaty. Environmental and Experimental Botany, 151, 55–63. 10.1016/j.envexpbot.2018.04.010

Heichel, G. H., & Anagnostakis, S. L. (1978). Stomatal Response to Light of *Solanum pennellii*, *Lycopersicon esculentum*, and a Graft-induced Chimera 1. Plant Physiology, 62(3), 387–390. doi:10.1104/pp.62.3.387

Hetherington, A. M., & Woodward, F. I. (2003). The role of stomata in sensing and driving environmental change. Nature, 424(6951), 901–908. doi:10.1038/nature01843

Ibanez, T., Ainsworth, A., Gross, J., Price, J. P., Webb, E. L., & Hart, P. J. (2021). Rarity patterns of woody plant species are associated with life form and diversification rates in Pacific islands forests. American Journal of Botany, 108(6), 946–957. 10.1002/ajb2.1687

Jin, Y., & Qian, H. (2019). V.PhyloMaker: an R package that can generate very large phylogenies for vascular plants. Ecography, 42(8), 1353–1359. 10.1111/ecog.04434

Jordan, G. J., Carpenter, R. J., & Brodribb, T. J. (2014). Using fossil leaves as evidence for open vegetation. Palaeogeography, Palaeoclimatology, Palaeoecology, 395, 168–175. 10.1016/j.palaeo.2013.12.035

Joswig, J. S., Wirth, C., Schuman, M. C., Kattge, J., Reu, B., Wright, I. J., . . . Mahecha, M. D. (2022). Climatic and soil factors explain the two-dimensional spectrum of global plant trait variation. Nature Ecology & Evolution, 6(1), 36–50. doi:10.1038/s41559-021-01616-8

Li, L., McCormack, M. L., Ma, C., Kong, D., Zhang, Q., Chen, X., . . . Guo, D. (2015). Leaf economics and hydraulic traits are decoupled in five species-rich tropical-subtropical forests. Ecology Letters, 18(9), 899–906. 10.1111/ele.12466

Liang, X., Xu, X., Wang, Z., He, L., Zhang, K., Liang, B., . . . Yang, W. (2022). StomataScorer: a portable and high-throughput leaf stomata trait scorer combined with deep learning and an improved CV model. Plant Biotechnology Journal, 20(3), 577–591. 10.1111/pbi.13741

Liu, C., He, N., Zhang, J., Li, Y., Wang, Q., Sack, L., & Yu, G. (2018). Variation of stomatal traits from cold temperate to tropical forests and association with water use efficiency. Functional Ecology, 32(1), 20–28. doi:10.1111/1365-2435.12973

Marek, S., Tomaszewski, D., Żytkowiak, R., Jasińska, A., Zadworny, M., Boratyńska, K., . . . Wyka, T. P. (2022). Stomatal density in Pinus sylvestris as an indicator of temperature rather than CO2: Evidence from a pan-European transect. Plant Cell and Environment, 45(1), 121–132. 10.1111/pce.14220

McKown, A. D., Guy, R. D., Quamme, L., Klápště, J., La Mantia, J., Constabel, C. P., . . . Azam, M. S. (2014). Association genetics, geography and ecophysiology link stomatal patterning in *Populus trichocarpa* with carbon gain and disease resistance trade-offs. Molecular Ecology, 23(23), 5771–5790. doi:10.1111/mec.12969

Melo, D., Porto, A., Cheverud, J. M., & Marroig, G. (2016). Modularity: Genes, Development, and Evolution. *Annual Review of Ecology*, Evolution, and Systematics, 47(1), 463–486. doi:10.1146/annurev-ecolsys-121415-032409

Milla, R., de Diego-Vico, N., & Martín-Robles, N. (2013). Shifts in stomatal traits following the domestication of plant species. Journal of Experimental Botany, 64(11), 3137–3146. doi:10.1093/jxb/ert147

Mott, K. A., Gibson, A. C., & O’leary, J. W. (1982). The adaptive significance of amphistomatic leaves. Plant, Cell & Environment, 5(6), 455–460. 10.1111/1365-3040.ep11611750

Muir, C. D. (2015). Making pore choices: repeated regime shifts in stomatal ratio. Proceedings of the Royal Society B: Biological Sciences, 282(1813), 20151498. doi:doi:10.1098/rspb.2015.1498

Muir, C. D. (2018). Light and growth form interact to shape stomatal ratio among British angiosperms. New Phytologist, 218(1), 242–252. 10.1111/nph.14956

Muir, C. D. (2019). Is Amphistomy an Adaptation to High Light? Optimality Models of Stomatal Traits along Light Gradients. Integrative and Comparative Biology, 59(3), 571–584. doi:10.1093/icb/icz085 %J Integrative and Comparative Biology

Münkemüller, T., Lavergne, S., Bzeznik, B., Dray, S., Jombart, T., Schiffers, K., & Thuiller, W. (2012). How to measure and test phylogenetic signal. Methods in Ecology and Evolution, 3(4), 743–756. 10.1111/j.2041-210X.2012.00196.x

Paradis, E., & Schliep, K. (2018). ape 5.0: an environment for modern phylogenetics and evolutionary analyses in R. Bioinformatics, 35(3), 526–528. doi:10.1093/bioinformatics/bty633

Parkhurst, D. F. (1978). The adaptive significance of stomatal occurrence on one or both surfaces of leaves. Journal of Ecology, 66(2), 367–383. doi:10.2307/2259142

Pearson, M., Davies, W. J., & Mansfield, T. A. (1995). Asymmetric responses of adaxial and abaxial stomata to elevated CO_2_: impacts on the control of gas exchange by leaves. Plant, Cell & Environment, 18(8), 837–843. doi:j.1365-3040.1995.tb00592.x

Raven, J. A. (2002). Selection pressures on stomatal evolution. New Phytologist, 153(3), 371–386. 10.1046/j.0028-646X.2001.00334.x

Revell, L. J. (2012). phytools: an R package for phylogenetic comparative biology (and other things). Methods in Ecology and Evolution, 3(2), 217–223. 10.1111/j.2041-210X.2011.00169.x

Richardson, F., Brodribb, T. J., & Jordan, G. J. (2017). Amphistomatic leaf surfaces independently regulate gas exchange in response to variations in evaporative demand. Tree Physiology, 37(7), 869–878. doi:10.1093/treephys/tpx073

Royer, D. L. (2001). Stomatal density and stomatal index as indicators of paleoatmospheric CO2 concentration. Review of Palaeobotany and Palynology, 114(1), 1–28. 10.1016/S0034-6667(00)00074-9

Sakoda, K., Watanabe, T., Sukemura, S., Kobayashi, S., Nagasaki, Y., Tanaka, Y., & Shiraiwa, T. (2019). Genetic diversity in stomatal density among soybeans elucidated using high-throughput technique based on an algorithm for object detection. Scientific Reports, 9(1), 7610. doi:10.1038/s41598-019-44127-0

Sardans, J., Vallicrosa, H., Zuccarini, P., Farré-Armengol, G., Fernández-Martínez, M., Peguero, G., . . . Peñuelas, J. (2021). Empirical support for the biogeochemical niche hypothesis in forest trees. Nature Ecology & Evolution, 5(2), 184–194. doi:10.1038/s41559-020-01348-1

Wall, S., Vialet-Chabrand, S., Davey, P., Van Rie, J., Galle, A., Cockram, J., & Lawson, T. (2022). Stomata on the abaxial and adaxial leaf surfaces contribute differently to leaf gas exchange and photosynthesis in wheat. New Phytologist, n/a(n/a). doi:10.1111/nph.18257

Xiong, D., & Flexas, J. (2020). From one side to two sides: the effects of stomatal distribution on photosynthesis. New Phytologist, 228(6), 1754–1766. 10.1111/nph.16801

Yan, Z., Sardans, J., Peñuelas, J., Detto, M., Smith, N. G., Wang, H., . . . Wu, J. (2023). Global patterns and drivers of leaf photosynthetic capacity: The relative importance of environmental factors and evolutionary history. Global Ecology and Biogeography, 32(5), 668–682. 10.1111/geb.13660

Yu, G., Smith, D. K., Zhu, H., Guan, Y., & Lam, T. T.-Y. (2017). ggtree: an r package for visualization and annotation of phylogenetic trees with their covariates and other associated data. Methods in Ecology and Evolution, 8(1), 28–36. 10.1111/2041-210X.12628

