## Supplementary material for "A continental-scale analysis reveals the latitudinal gradient of stomatal density across amphistomatous species: Evolutionary history vs. present-day environment": SI

Running title：Stomatal density of amphistomatous species

Congcong Liu^1,2,3#*^, Kexiang Huang^1,2#^, Yifei Zhao^1,2^, Ying Li^3^, and Nianpeng He^3,4,5*^

^1^ Key Laboratory of Ecology and Environment in Minority Areas (Minzu University of China), National Ethnic Affairs Commission, Beijing 100081, China

^2^ College of Life and Environmental Sciences, Minzu University of China, Beijing 100081, China

^3^ Key Laboratory of Ecosystem Network Observation and Modeling, Institute of Geographic Sciences and Natural Resources Research, Chinese Academy of Sciences, Beijing 100101, China

^4^ Center for Ecological Research, Northeast Forestry University, Harbin 150040, China

^5^ Earth Critical Zone and Flux Research Station of Xing’an Mountains, Chinese Academy of Sciences, Daxing'anling 165200, China

^#^ These authors contributed equally.

ORCID ID

Nianpeng He 0000-0002-0458-5953

Congcong Liu 0000-0003-3949-4194

**Table S1 Overview of the data used in this study.**

|  | Species-site combinations with coordinates | Species-site combinations without coordinates | Total |
| --- | --- | --- | --- |
| Woody species | 58 | 10 | 68 |
| Herbaceous species | 360 | 58 | 418 |
| Total | 418 | 68 | 486 |

**Table S2 Summary of environmental variables used in this study.**

| Full name | Abbr. | Unit | Note |
| --- | --- | --- | --- |
| Aridity index (MAP/PET) | AI | Unitless |  |
| Mean annual temperature | MAT | °C |  |
| Temperature seasonality | TS | Unitless | standard deviation ×100 |
| Mean annual precipitation | MAP | mm |  |
| Precipitation seasonality | PS | Unitless | Coefficient of Variation |
| Solar radiation | Solar | kJ m^-2^ day^-1^ |  |
| Soil nitrogen content | N | % of weight |  |
| Soil sand content | SAND | % of weight |  |
| Soil silt content | SILT | % of weight |  |
| Soil bulk density | BD | g mm^-3^ |  |
| Soil pH | pH | Unitless |  |

**Table S3 Changes in stomatal traits of all species, woody species and herbaceous species.**

|  | Mean ± SE |
| --- | --- |
| **All species** |  |
| SD_ad_ | 135.82 ± 8.25 |
| SD_ab_ | 174.59 ± 8.58 |
| SD_total_ | 310.41 ± 15.16 |
| SR | 0.95 ± 0.06 |
| **Wood species** |  |
| SD_ad_ | 229.34 ± 58.60 |
| SD_ab_ | 188.09 ± 42.42 |
| SD_total_ | 417.43 ± 99.47 |
| SR | 1.62 ± 0.38 |
| **Herbaceous species** |  |
| SD_ad_ | 127.42 ± 7.14 |
| SD_ab_ | 173.37 ± 8.57 |
| SD_total_ | 300.79 ± 13.85 |
| SR | 0.89 ± 0.05 |

SD_ad_, adaxial stomatal density; SD_ab_, abaxial stomatal density; SD_total_; total stomatal density; SR, stomatal ratio.

**Table S4 Summary of the effects of growth form on the adaxial stomatal density (SD_ad_) across amphistomatous species.**

| **Bayesian model** | **Model statistics** | | |  |  |  |
| --- | --- | --- | --- | --- | --- | --- |
|  | **The statistics of fixed variables** | | |  |  |  |
| MCMCglmm.updateable  (SD_ad_ ~growth, random=~ phylogeny + species) | Variable | Post.mean | Lower 95% CI | Upper 95% CI | Eff.samp | pMCMC |
|  | Intercept | 125.71 | 72.46 | 175.63 | 1700 | < 0.001 |
|  | GrowthWoody | 87.24 | 27.87 | 142.98 | 1700 | 0.004 |
|  | **R^2^ of the model** | | |  |  |  |
|  | R^2^m = 0.044; R^2^c = 0.708 | | | | | |

The linear regression is estimated using the Bayesian phylogenetic linear mixed model with phylogeny and species as random factors. Post.mean, posterior mean; Eff.samp, the effective sample size; pMCMC, p-value from Monte Carlo sampling by Markov Chain. Lower 95% CI, the lower limit of the 95% confidence interval for slope. Upper 95% CI, the upper limit of the 95% confidence interval for slope. R^2^m, marginal R^2^ (fixed effects only); R^2^c, conditional R^2^ (both fixed and random effects).

**Table S5 Summary of the effects of growth form on the abaxial stomatal density (SD_ab_) across amphistomatous species.**

| **Bayesian model** | **Model statistics** | | |  |  |  |
| --- | --- | --- | --- | --- | --- | --- |
|  | **The statistics of fixed variables** | | |  |  |  |
| MCMCglmm.updateable  (SD_ab_ ~growth, random=~ phylogeny + species) | Variable | Post.mean | Lower 95% CI | Upper 95% CI | Eff.samp | pMCMC |
|  | Intercept | 160.03 | 53.85 | 284.14 | 1612 | 0.009 |
|  | GrowthWoody | 29.61 | -42.94 | 99.16 | 1700 | 0.4153 |
|  | **R^2^ of the model** | | |  |  |  |
|  | R^2^m =0.003; R^2^c = 0.734 | | | | | |

The linear regression is estimated using the Bayesian phylogenetic linear mixed model with phylogeny and species as random factors. Post.mean, posterior mean; Eff.samp, the effective sample size; pMCMC, p-value from Monte Carlo sampling by Markov Chain. Lower 95% CI, the lower limit of the 95% confidence interval for slope. Upper 95% CI, the upper limit of the 95% confidence interval for slope. R^2^m, marginal R^2^ (fixed effects only); R^2^c, conditional R^2^ (both fixed and random effects).

**Table S6 Summary of the effects of growth form on the total stomatal density (SD_total_) across amphistomatous species.**

| **Bayesian model** | **Model statistics** | | |  |  |  |
| --- | --- | --- | --- | --- | --- | --- |
|  | **The statistics of fixed variables** | | |  |  |  |
| MCMCglmm.updateable  (SD_total_ ~growth, random=~ phylogeny + species) | Variable | Post.mean | Lower 95% CI | Upper 95% CI | Eff.samp | pMCMC |
|  | Intercept | 293.675 | 180.408 | 399.788 | 1373 | <0.001 |
|  | GrowthWoody | 102.209 | -5.169 | 209.624 | 1700 | 0.0565 |
|  | **R^2^ of the model** | | |  |  |  |
|  | R^2^m =0.017; R^2^c = 0.646 | | | | | |

The linear regression is estimated using the Bayesian phylogenetic linear mixed model with phylogeny and species as random factors. Post.mean, posterior mean; Eff.samp, the effective sample size; pMCMC, p-value from Monte Carlo sampling by Markov Chain. Lower 95% CI, the lower limit of the 95% confidence interval for slope. Upper 95% CI, the upper limit of the 95% confidence interval for slope. R^2^m, marginal R^2^ (fixed effects only); R^2^c, conditional R^2^ (both fixed and random effects).

**Table S7 Summary of the effects of growth form on the stomatal ratio (SR) across amphistomatous species.**

| **Bayesian model** | **Model statistics** | | |  |  |  |
| --- | --- | --- | --- | --- | --- | --- |
|  | **The statistics of fixed variables** | | |  |  |  |
| MCMCglmm.updateable  (SR ~growth, random=~ phylogeny + species) | Variable | Post.mean | Lower 95% CI | Upper 95% CI | Eff.samp | pMCMC |
|  | Intercept | 0.9764 | 0.0826 | 1.8674 | 1858 | 0.04 |
|  | GrowthWoody | 0.5006 | -0.0060 | 0.9878 | 1700 | 0.0447 |
|  | **R^2^ of the model** | | |  |  |  |
|  | R^2^m =0.014; R^2^c = 0.667 | | | | | |

The linear regression is estimated using the Bayesian phylogenetic linear mixed model with phylogeny and species as random factors. Post.mean, posterior mean; Eff.samp, the effective sample size; pMCMC, p-value from Monte Carlo sampling by Markov Chain. Lower 95% CI, the lower limit of the 95% confidence interval for slope. Upper 95% CI, the upper limit of the 95% confidence interval for slope. R^2^m, marginal R^2^ (fixed effects only); R^2^c, conditional R^2^ (both fixed and random effects).

**Table S8 Relationships between the adaxial stomatal density (SD_ad_) and abaxial stomatal density (SD_ab_).**

| **Bayesian model** | **Model statistics** | | |  |  |  |
| --- | --- | --- | --- | --- | --- | --- |
|  | **The statistics of fixed variables** | | |  |  |  |
| MCMCglmm.updateable  (log_10_(SD_ad_) ~ log_10_(SD_ab_), random=~ phylogeny + species) | Variable | Post.mean | Lower 95% CI | Upper 95% CI | Eff.samp | pMCMC |
|  | Intercept | 0.3414 | -0.0562 | 0.6867 | 2258 | 0.0847 |
|  | log_10_(SD_ab_) | 0.7591 | 0.6819 | 0.8368 | 1700 | < 0.001 |
|  | **R^2^ of the model** | | |  |  |  |
|  | R^2^m = 0.242; R^2^c =0.875 | | | | | |

The linear regression is estimated using the Bayesian phylogenetic linear mixed model with phylogeny and species as random factors. Post.mean, posterior mean; Eff.samp, the effective sample size; pMCMC, p-value from Monte Carlo sampling by Markov Chain. Lower 95% CI, the lower limit of the 95% confidence interval for slope. Upper 95% CI, the upper limit of the 95% confidence interval for slope. R^2^m, marginal R^2^ (fixed effects only); R^2^c, conditional R^2^ (both fixed and random effects).

**Table S9 The influence of the adaxial stomatal density (SD_ad_) and abaxial stomatal density (SD_ab_) on stomatal ratio (B) across amphistomatous species.**

| **Bayesian model** | **Model statistics** | | |  |  |  |
| --- | --- | --- | --- | --- | --- | --- |
|  | **The statistics of fixed variables** | | |  |  |  |
| MCMCglmm.updateable  (scale(SD_ad_) ~ scale(SD_ab_)+ scale(SD_ab_), random=~ phylogeny + species) | Variable | Post.mean | Lower 95% CI | Upper 95% CI | Eff.samp | pMCMC |
|  | Intercept | 0.0095 | -0.5035 | 0.5854 | 1700 | 0.969 |
|  | scale (SD_ad_) | 0.5697 | 0.4859 | 0.6673 | 1700 | <0.001 |
|  | scale (SD_ab_) | -0.6008 | -0.6964 | -0.5198 | 1700 | <0.001 |
|  | **R^2^ of the model** | | |  |  |  |
|  | R^2^m = 0.244; R^2^c = 0.688 | | | | | |

The linear regression is estimated using the Bayesian phylogenetic linear mixed model with phylogeny and species as random factors. Post.mean, posterior mean; Eff.samp, the effective sample size; pMCMC, p-value from Monte Carlo sampling by Markov Chain. Lower 95% CI, the lower limit of the 95% confidence interval for slope. Upper 95% CI, the upper limit of the 95% confidence interval for slope. R^2^m, marginal R^2^ (fixed effects only); R^2^c, conditional R^2^ (both fixed and random effects).

**Table S10 Relationships between the adaxial stomatal density (SD_ad_) and latitude across amphistomatous species.**

| **Bayesian model** | **Model statistics** | | |  |  |  |
| --- | --- | --- | --- | --- | --- | --- |
|  | **The statistics of fixed variables** | | |  |  |  |
| MCMCglmm.updateable  (SD_ad_ ~latitude, random=~ phylogeny + species) | Variable | Post.mean | Lower 95% CI | Upper 95% CI | Eff.samp | pMCMC |
|  | Intercept | 254.339 | 161.786 | 362.756 | 1700 | <0.001 |
|  | latitude | -4.088 | -6.039 | -2.204 | 1700 | <0.001 |
|  | **R^2^ of the model** | | |  |  |  |
|  | R^2^m = 0.040; R^2^c = 0.738 | | | | | |

The linear regression is estimated using the Bayesian phylogenetic linear mixed model with phylogeny and species as random factors. Post.mean, posterior mean; Eff.samp, the effective sample size; pMCMC, p-value from Monte Carlo sampling by Markov Chain. Lower 95% CI, the lower limit of the 95% confidence interval for slope. Upper 95% CI, the upper limit of the 95% confidence interval for slope. R^2^m, marginal R^2^ (fixed effects only); R^2^c, conditional R^2^ (both fixed and random effects).

**Table S11 Relationships between the abaxial stomatal density (SD_ab_) and latitude across amphistomatous species.**

| **Bayesian model** | **Model statistics** | | |  |  |  |
| --- | --- | --- | --- | --- | --- | --- |
|  | **The statistics of fixed variables** | | |  |  |  |
| MCMCglmm.updateable  (SD_ab_ ~latitude, random=~ phylogeny + species) | Variable | Post.mean | Lower 95% CI | Upper 95% CI | Eff.samp | pMCMC |
|  | Intercept | 344.074 | 226.931 | 473.655 | 1700 | <0.001 |
|  | latitude | -5.929 | -8.074 | -3.602 | 1474 | <0.001 |
|  | **R^2^ of the model** | | |  |  |  |
|  | R^2^m = 0.056; R^2^c = 0.763 | | | | | |

The linear regression is estimated using the Bayesian phylogenetic linear mixed model with phylogeny and species as random factors. Post.mean, posterior mean; Eff.samp, the effective sample size; pMCMC, p-value from Monte Carlo sampling by Markov Chain. Lower 95% CI, the lower limit of the 95% confidence interval for slope. Upper 95% CI, the upper limit of the 95% confidence interval for slope. R^2^m, marginal R^2^ (fixed effects only); R^2^c, conditional R^2^ (both fixed and random effects).

**Table S12 Relationships between the total stomatal density (SD_total_) and latitude across amphistomatous species.**

| **Bayesian model** | **Model statistics** | | |  |  |  |
| --- | --- | --- | --- | --- | --- | --- |
|  | **The statistics of fixed variables** | | |  |  |  |
| MCMCglmm.updateable  (SD_total_ ~latitude, random=~ phylogeny + species) | Variable | Post.mean | Lower 95% CI | Upper 95% CI | Eff.samp | pMCMC |
|  | Intercept | 596.225 | 394.621 | 789.289 | 1700 | <0.001 |
|  | latitude | -10.064 | -13.672 | -6.471 | 1700 | <0.001 |
|  | **R^2^ of the model** | | |  |  |  |
|  | R^2^m = 0.062; R^2^c = 0.741 | | | | | |

The linear regression is estimated using the Bayesian phylogenetic linear mixed model with phylogeny and species as random factors. Post.mean, posterior mean; Eff.samp, the effective sample size; pMCMC, p-value from Monte Carlo sampling by Markov Chain. Lower 95% CI, the lower limit of the 95% confidence interval for slope. Upper 95% CI, the upper limit of the 95% confidence interval for slope. R^2^m, marginal R^2^ (fixed effects only); R^2^c, conditional R^2^ (both fixed and random effects).

**Table S13 Relationships between the stomatal ratio (SR) and latitude across amphistomatous species.**

| **Bayesian model** | **Model statistics** | | |  |  |  |
| --- | --- | --- | --- | --- | --- | --- |
|  | **The statistics of fixed variables** | | |  |  |  |
| MCMCglmm.updateable  (SD_total_ ~latitude, random=~ phylogeny + species) | Variable | Post.mean | Lower 95% CI | Upper 95% CI | Eff.samp | pMCMC |
|  | Intercept | 0.0035 | -1.1040 | 1.2188 | 1526 | 0.9953 |
|  | latitude | 0.0312 | 0.0081 | 0.0503 | 1700 | 0.0047 |
|  | **R^2^ of the model** | | |  |  |  |
|  | R^2^m = 0.018; R^2^c = 0.699 | | | | | |

The linear regression is estimated using the Bayesian phylogenetic linear mixed model with phylogeny and species as random factors. Post.mean, posterior mean; Eff.samp, the effective sample size; pMCMC, p-value from Monte Carlo sampling by Markov Chain. Lower 95% CI, the lower limit of the 95% confidence interval for slope. Upper 95% CI, the upper limit of the 95% confidence interval for slope. R^2^m, marginal R^2^ (fixed effects only); R^2^c, conditional R^2^ (both fixed and random effects).

**Table S14** **Bivariate relationships between the adaxial stomatal density (SD_ad_) and environmental factors across amphistomatous species.**

| Variable | Post.mean | Lower 95% CI | Upper 95% CI | Eff.samp | pMCMC | R^2^ of the model |
| --- | --- | --- | --- | --- | --- | --- |
| Intercept | 96.7711 | 16.7073 | 167.3084 | 1530.58 | 0.0129 | R^2^m = 0.0057 |
| AI | 24.1578 | -12.8912 | 60.3601 | 1700 | 0.1906 | R^2^c = 0.7083 |
| Intercept | 99.5685 | 27.3908 | 175.5780 | 1700 | 0.0094 | R^2^m = 0.0030 |
| MAT | 1.0585 | -0.8891 | 2.8788 | 1835.306 | 0.2871 | R^2^c = 0.6951 |
| **Intercept** | **188.0981** | **105.3255** | **272.6198** | **1700** | **0.0006** | **R^2^m = 0.0341** |
| **TS** | **-0.8714** | **-1.3124** | **-0.4293** | **1700** | **0.0006** | **R^2^c = 0.7072** |
| Intercept | 94.5296 | 17.0260 | 168.6880 | 1854.364 | 0.0153 | R^2^m = 0.0049 |
| MAP | 0.0224 | -0.0088 | 0.0504 | 1700 | 0.1329 | R^2^c = 0.7082 |
| Intercept | 76.8382 | -2.3327 | 158.9554 | 1650.93 | 0.0659 | R^2^m = 0.0071 |
| PS | 0.4053 | -0.0067 | 0.8569 | 1700 | 0.0600 | R^2^c = 0.6944 |
| Intercept | 19.9635 | -141.6320 | 185.5050 | 1700 | 0.8271 | R^2^m = 0.0028 |
| Solar | 0.0059 | -0.0045 | 0.0152 | 1276.565 | 0.2506 | R^2^c = 0.6888 |
| Intercept | 108.6105 | 26.2938 | 178.9749 | 1700 | 0.0071 | R^2^m = 0.0000 |
| N | 4.6722 | -52.2369 | 68.3798 | 1820.046 | 0.8976 | R^2^c = 0.7045 |
| Intercept | 120.4948 | 44.8576 | 192.7485 | 1700 | 0.0059 | R^2^m = 0.0015 |
| Sand | -0.3064 | -0.8867 | 0.1634 | 1700 | 0.2565 | R^2^c = 0.6949 |
| Intercept | 95.5124 | 18.0327 | 183.9283 | 1486.246 | 0.0294 | R^2^m = 0.0009 |
| Silt | 0.3364 | -0.4073 | 1.0137 | 1823.018 | 0.3671 | R^2^c = 0.6977 |
| Intercept | 107.6140 | -17.2894 | 219.9863 | 1700 | 0.0753 | R^2^m = 0.0000 |
| BD | 0.3495 | -75.7381 | 80.7374 | 1700 | 0.9847 | R^2^c = 0.6959 |
| Intercept | 127.1770 | 19.2332 | 234.5519 | 1700 | 0.0212 | R^2^m = 0.0003 |
| pH | -2.1529 | -12.0915 | 9.2152 | 1502.148 | 0.7024 | R^2^c = 0.7069 |

The linear regressions were estimated using the Bayesian phylogenetic linear mixed model with phylogeny and species as random factors: MCMCglmm.updateable (SD_ad_ ~environment, random=~ phylogeny + species). The abbreviations of environmental factors were shown in Table S1. Significant bivariate relationships are indicated in bold. Post.mean, posterior mean; Eff.samp, the effective sample size; pMCMC, p-value from Monte Carlo sampling by Markov Chain. Lower 95% CI, the lower limit of the 95% confidence interval for slope. Upper 95% CI, the upper limit of the 95% confidence interval for slope. R^2^m, marginal R^2^ (fixed effects only); R^2^c, conditional R^2^ (both fixed and random effects).

Bold values indicate that the 95% credible intervals exclude zero and, therefore, are statistically significant.

**Table S15 Bivariate relationships between the abaxial stomatal density (SD_ab_) and environmental factors across amphistomatous species.**

| Variable | Post.mean | Lower 95% CI | Upper 95% CI | Eff.samp | pMCMC | R^2^ of the model |
| --- | --- | --- | --- | --- | --- | --- |
| **Intercept** | **94.6568** | **-12.1436** | **203.8639** | **1700** | **0.0976** | **R^2^m = 0.0271** |
| **AI** | **70.9273** | **27.3296** | **113.6404** | **1848.125** | **0.0012** | **R^2^c = 0.7758** |
| **Intercept** | **113.4229** | **13.0859** | **208.9725** | **1776.439** | **0.0259** | **R^2^m = 0.0113** |
| **MAT** | **2.6780** | **0.3704** | **4.8809** | **1700** | **0.0200** | **R^2^c = 0.7468** |
| **Intercept** | **258.5391** | **143.9161** | **362.5732** | **2140.953** | **0.0006** | **R^2^m = 0.0499** |
| **TS** | **-1.3833** | **-1.9590** | **-0.8395** | **1700** | **0.0006** | **R^2^c = 0.7742** |
| **Intercept** | **107.7488** | **-0.2807** | **205.5671** | **1700** | **0.0529** | **R^2^m = 0.0133** |
| **MAP** | **0.0481** | **0.0126** | **0.0847** | **1700** | **0.0106** | **R^2^c = 0.7562** |
| Intercept | 134.5931 | 21.7162 | 228.8639 | 1906.321 | 0.0118 | R^2^m = 0.0000 |
| PS | -0.0110 | -0.5570 | 0.4972 | 2165.687 | 0.9647 | R^2^c = 0.7544 |
| Intercept | 140.9505 | -52.9283 | 365.2232 | 1700 | 0.1788 | R^2^m = 0.0000 |
| Solar | -0.0004 | -0.0128 | 0.0118 | 1700 | 0.9588 | R^2^c = 0.7602 |
| Intercept | 129.4931 | 19.5105 | 234.4284 | 1700 | 0.0188 | R^2^m = 0.0003 |
| N | 18.0740 | -46.1601 | 85.9835 | 1700 | 0.6024 | R^2^c = 0.7598 |
| Intercept | 135.5255 | 33.2616 | 243.4002 | 2057.458 | 0.0129 | R^2^m = 0.0000 |
| Sand | 0.0064 | -0.6320 | 0.6551 | 1572.022 | 0.9588 | R^2^c = 0.7633 |
| Intercept | 123.6648 | 8.8892 | 238.0736 | 1700 | 0.0329 | R^2^m = 0.0004 |
| Silt | 0.3068 | -0.6417 | 1.1598 | 1700 | 0.5000 | R^2^c = 0.7641 |
| Intercept | 189.6267 | 34.9588 | 347.0554 | 1700 | 0.0141 | R^2^m = 0.0008 |
| BD | -44.2574 | -133.2084 | 52.2202 | 1700 | 0.3588 | R^2^c = 0.7627 |
| Intercept | 219.2322 | 93.5034 | 369.3594 | 1700 | 0.0024 | R^2^m = 0.0054 |
| pH | -11.2154 | -23.4944 | 0.7977 | 1700 | 0.0788 | R^2^c = 0.7634 |

The linear regressions were estimated using the Bayesian phylogenetic linear mixed model with phylogeny and species as random factors: MCMCglmm.updateable (SD_ab_ ~environment, random=~ phylogeny + species). The abbreviations of environmental factors were shown in Table S1. Significant bivariate relationships are indicated in bold. Post.mean, posterior mean; Eff.samp, the effective sample size; pMCMC, p-value from Monte Carlo sampling by Markov Chain. Lower 95% CI, the lower limit of the 95% confidence interval for slope. Upper 95% CI, the upper limit of the 95% confidence interval for slope. R^2^m, marginal R^2^ (fixed effects only); R^2^c, conditional R^2^ (both fixed and random effects).

Bold values indicate that the 95% credible intervals exclude zero and, therefore, are statistically significant.

**Table S16 Bivariate relationships between the total stomatal density (SD_total_) and environmental factors across amphistomatous species.**

| Variable | Post.mean | Lower 95% CI | Upper 95% CI | Eff.samp | pMCMC | R^2^ of the model |
| --- | --- | --- | --- | --- | --- | --- |
| **Intercept** | **190.4434** | **18.8835** | **350.4296** | **1700** | **0.0282** | **R^2^m = 0.0207** |
| **AI** | **97.7158** | **29.3326** | **172.5815** | **1700** | **0.0141** | **R^2^c = 0.7459** |
| **Intercept** | **213.6401** | **58.3515** | **371.0825** | **1667.1** | **0.0094** | **R^2^m = 0.0088** |
| **MAT** | **3.6893** | **0.2288** | **7.4265** | **1366.25** | **0.0494** | **R^2^c = 0.7104** |
| **Intercept** | **445.6900** | **273.9214** | **602.5420** | **1700** | **0.0006** | **R^2^m = 0.0528** |
| **TS** | **-2.2411** | **-3.1576** | **-1.3905** | **2556.816** | **0.0006** | **R^2^c = 0.7375** |
| **Intercept** | **202.2750** | **8.9046** | **350.7666** | **1700** | **0.0259** | **R^2^m = 0.0107** |
| **MAP** | **0.0707** | **0.0165** | **0.1293** | **1700** | **0.0176** | **R^2^c = 0.7395** |
| Intercept | 212.9431 | 40.3396 | 371.4804 | 1700 | 0.0176 | R^2^m = 0.0017 |
| PS | 0.4056 | -0.3992 | 1.3283 | 1700 | 0.3424 | R^2^c = 0.7108 |
| Intercept | 169.7141 | -153.1498 | 474.7440 | 1700 | 0.2988 | R^2^m = 0.0005 |
| Solar | 0.0050 | -0.0150 | 0.0228 | 1700 | 0.6035 | R^2^c = 0.7124 |
| Intercept | 240.1383 | 54.9764 | 387.9377 | 1700 | 0.0047 | R^2^m = 0.0002 |
| N | 24.9753 | -83.1840 | 135.2241 | 1900.342 | 0.6647 | R^2^c = 0.7195 |
| Intercept | 260.4519 | 94.0028 | 430.4963 | 1700 | 0.0024 | R^2^m = 0.0005 |
| Sand | -0.3587 | -1.4561 | 0.6771 | 1700 | 0.5106 | R^2^c = 0.7252 |
| Intercept | 215.9887 | 36.8943 | 378.3573 | 1700 | 0.0200 | R^2^m = 0.0009 |
| Silt | 0.7129 | -0.7783 | 2.0645 | 1700 | 0.3106 | R^2^c = 0.7238 |
| Intercept | 290.6915 | 41.9846 | 526.0618 | 1700 | 0.0176 | R^2^m = 0.0002 |
| BD | -37.0152 | -202.1713 | 109.8658 | 1700 | 0.6424 | R^2^c = 0.7247 |
| Intercept | 340.0364 | 122.0587 | 538.1708 | 1700 | 0.0047 | R^2^m = 0.0030 |
| pH | -13.0759 | -31.5718 | 8.9871 | 1700 | 0.2165 | R^2^c = 0.7273 |

The linear regressions were estimated using the Bayesian phylogenetic linear mixed model with phylogeny and species as random factors: MCMCglmm.updateable (SD_total_ ~environment, random=~ phylogeny + species). The abbreviations of environmental factors were shown in Table S1. Significant bivariate relationships are indicated in bold. Post.mean, posterior mean; Eff.samp, the effective sample size; pMCMC, p-value from Monte Carlo sampling by Markov Chain. Lower 95% CI, the lower limit of the 95% confidence interval for slope. Upper 95% CI, the upper limit of the 95% confidence interval for slope. R^2^m, marginal R^2^ (fixed effects only); R^2^c, conditional R^2^ (both fixed and random effects).

Bold values indicate that the 95% credible intervals exclude zero and, therefore, are statistically significant.

**Table S17** **Bivariate relationships between the stomatal ratio (SR) and environmental factors across amphistomatous species.**

| Variable | Post.mean | Lower 95% CI | Upper 95% CI | Eff.samp | pMCMC | R^2^ of the model |
| --- | --- | --- | --- | --- | --- | --- |
| Intercept | 1.1680 | 0.1928 | 2.1194 | 2076.013 | 0.0176 | R^2^m = 0.0024 |
| AI | -0.1821 | -0.5869 | 0.2278 | 1872.96 | 0.3694 | R^2^c = 0.6755 |
| Intercept | 1.1326 | 0.2861 | 1.9738 | 1700 | 0.0165 | R^2^m = 0.0006 |
| MAT | -0.0055 | -0.0276 | 0.0138 | 1700 | 0.5859 | R^2^c = 0.6788 |
| **Intercept** | **0.4655** | **-0.5974** | **1.4349** | **1700** | **0.3718** | **R^2^m = 0.0154** |
| **TS** | **0.0070** | **0.0020** | **0.0128** | **1866.446** | **0.0118** | **R^2^c = 0.6870** |
| Intercept | 1.1631 | 0.2646 | 2.0744 | 1700 | 0.0153 | R^2^m = 0.0010 |
| MAP | -0.0001 | -0.0004 | 0.0002 | 1700 | 0.4941 | R^2^c = 0.6794 |
| Intercept | 0.8004 | -0.1126 | 1.8386 | 1700 | 0.1047 | R^2^m = 0.0038 |
| PS | 0.0035 | -0.0015 | 0.0083 | 1700 | 0.1647 | R^2^c = 0.6737 |
| Intercept | 1.7409 | -0.1604 | 3.5004 | 1370.428 | 0.0682 | R^2^m = 0.0011 |
| Solar | 0.0000 | -0.0002 | 0.0001 | 1547.034 | 0.4235 | R^2^c = 0.6820 |
| Intercept | 1.0899 | 0.2247 | 1.9041 | 1700 | 0.0153 | R^2^m = 0.0000 |
| N | 0.0419 | -0.6543 | 0.6602 | 1700 | 0.9035 | R^2^c = 0.6773 |
| Intercept | 1.1949 | 0.2933 | 2.2179 | 1524.277 | 0.0153 | R^2^m = 0.0010 |
| Sand | -0.0029 | -0.0095 | 0.0032 | 1700 | 0.3635 | R^2^c = 0.6769 |
| Intercept | 1.0237 | 0.0658 | 1.9663 | 1700 | 0.0353 | R^2^m = 0.0001 |
| Silt | 0.0014 | -0.0070 | 0.0098 | 1700 | 0.7353 | R^2^c = 0.6771 |
| Intercept | 0.4808 | -0.9774 | 1.9227 | 1579.722 | 0.5176 | R^2^m = 0.0013 |
| BD | 0.4971 | -0.4231 | 1.4963 | 1700 | 0.2718 | R^2^c = 0.6769 |
| Intercept | 0.6853 | -0.5069 | 1.8717 | 1582.458 | 0.2671 | R^2^m = 0.0017 |
| pH | 0.0562 | -0.0713 | 0.1666 | 1700 | 0.3612 | R^2^c = 0.6748 |

The linear regressions were estimated using the Bayesian phylogenetic linear mixed model with phylogeny and species as random factors: MCMCglmm.updateable (SR ~environment, random=~ phylogeny + species). The abbreviations of environmental factors were shown in Table S1. Significant bivariate relationships are indicated in bold. Post.mean, posterior mean; Eff.samp, the effective sample size; pMCMC, p-value from Monte Carlo sampling by Markov Chain. Lower 95% CI, the lower limit of the 95% confidence interval for slope. Upper 95% CI, the upper limit of the 95% confidence interval for slope. R^2^m, marginal R^2^ (fixed effects only); R^2^c, conditional R^2^ (both fixed and random effects).

Bold values indicate that the 95% credible intervals exclude zero and, therefore, are statistically significant.

**Table S18 Results from Bayesian phylogenetic linear mixed model of the adaxial stomatal density (SD_ad_), with fixed factors (i.e., environmental factors) and random factors (i.e., phylogeny + species) taken into account.**

| Bayesian model | Variable | Post.mean | Lower 95% CI | Upper 95% CI | Eff.samp | pMCMC | R^2^ of the model |
| --- | --- | --- | --- | --- | --- | --- | --- |
| MCMCglmm.updateable(SD_ad_~AI+MAT+TS+MAP+PS+solar+N+Sand+Silt+BD+pH, random=~ phylogeny + species) | (Intercept) | -0.0995 | -0.7780 | 0.5976 | 1895 | 0.7518 | R^2^m = 0.039 |
|  | AI | 0.0365 | -0.2686 | 0.3797 | 1700 | 0.8412 | R^2^p = 0.529 |
|  | MAT | 0.0735 | -0.0794 | 0.2215 | 1700 | 0.3518 | R^2^s = 0.098 |
|  | **TS** | **-0.2515** | **-0.4177** | **-0.0858** | **1430** | **0.0071** | R^2^c = 0.666 |
|  | MAP | -0.0183 | -0.2345 | 0.1970 | 1700 | 0.8224 |  |
|  | PS | 0.0275 | -0.1099 | 0.1480 | 1700 | 0.6577 |  |
|  | Solar | 0.0604 | -0.0964 | 0.1987 | 1275 | 0.4188 |  |
|  | N | 0.0683 | -0.0688 | 0.2002 | 1700 | 0.3259 |  |
|  | Sand | -0.0812 | -0.2908 | 0.1454 | 1700 | 0.4788 |  |
|  | Silt | -0.0143 | -0.2245 | 0.2185 | 1700 | 0.9035 |  |
|  | BD | 0.0532 | -0.0672 | 0.1669 | 1844 | 0.3659 |  |
|  | pH | 0.1205 | -0.0705 | 0.3016 | 1700 | 0.2424 |  |

The abbreviations of environmental factors were shown in Table S1. Significant effects of environmental factors on stomatal traits are indicated in bold. Post.mean, posterior mean; Eff.samp, the effective sample size; pMCMC, p-value from Monte Carlo sampling by Markov Chain. Lower 95% CI, the lower limit of the 95% confidence interval for slope. Upper 95% CI, the upper limit of the 95% confidence interval for slope. R^2^c, percentage of variance explained by all the model (fixed + random); R^2^m, percentage of variance explained by fixed factors; R^2^p, percentage of variance explained by phylogeny; R^2^s, percentage of variance explained by species.

Bold values indicate that the 95% credible intervals exclude zero and, therefore, are statistically significant.

**Table S19 Results from Bayesian phylogenetic linear mixed model of the abaxial stomatal density (SD_ab_), with fixed factors (i.e., environmental factors) and random factors (i.e., phylogeny + species) taken into account.**

| Bayesian model | Variable | Post.mean | Lower 95% CI | Upper 95% CI | Eff.samp | pMCMC | R^2^ of the model |
| --- | --- | --- | --- | --- | --- | --- | --- |
| MCMCglmm.updateable(SD_ab_~AI+MAT+TS+MAP+PS+solar+N+Sand+Silt+BD+pH, random=~ phylogeny + species) | (Intercept) | -0.2146 | -1.0372 | 0.6452 | 1576 | 0.6329 | R^2^m = 0.068 |
|  | AI | 0.1304 | -0.1559 | 0.4445 | 1465 | 0.4082 | R^2^p = 0.652 |
|  | MAT | 0.1335 | -0.0139 | 0.2881 | 1700 | 0.0800 | R^2^s = 0.060 |
|  | **TS** | **-0.3087** | **-0.4937** | **-0.1458** | **1852** | **0.0012** | R^2^c = 0.779 |
|  | MAP | -0.0100 | -0.2073 | 0.1985 | 1700 | 0.9541 |  |
|  | PS | -0.0800 | -0.2050 | 0.0432 | 1700 | 0.2188 |  |
|  | Solar | 0.0256 | -0.1294 | 0.1532 | 1700 | 0.7165 |  |
|  | N | 0.0527 | -0.0631 | 0.2024 | 1700 | 0.4459 |  |
|  | Sand | 0.1782 | -0.0352 | 0.3841 | 1700 | 0.0918 |  |
|  | Silt | 0.1853 | -0.0229 | 0.3938 | 1700 | 0.0788 |  |
|  | BD | -0.0279 | -0.1373 | 0.0897 | 1700 | 0.6247 |  |
|  | pH | 0.1349 | -0.0486 | 0.3237 | 1700 | 0.1647 |  |

The abbreviations of environmental factors were shown in Table S1. Significant effects of environmental factors on stomatal traits are indicated in bold. Post.mean, posterior mean; Eff.samp, the effective sample size; pMCMC, p-value from Monte Carlo sampling by Markov Chain. Lower 95% CI, the lower limit of the 95% confidence interval for slope. Upper 95% CI, the upper limit of the 95% confidence interval for slope. R^2^c, percentage of variance explained by all the model (fixed + random); R^2^m, percentage of variance explained by fixed factors; R^2^p, percentage of variance explained by phylogeny; R^2^s, percentage of variance explained by species.

Bold values indicate that the 95% credible intervals exclude zero and, therefore, are statistically significant.

**Table S20 Results from Bayesian phylogenetic linear mixed model of the total stomatal density (SD_total_), with fixed factors (i.e., environmental factors) and random factors (i.e., phylogeny + species) taken into account.**

| Bayesian model | Variable | Post.mean | Lower 95% CI | Upper 95% CI | Eff.samp | pMCMC | R^2^ of the model |
| --- | --- | --- | --- | --- | --- | --- | --- |
| MCMCglmm.updateable(SD_total_~AI+MAT+TS+MAP+PS+solar+N+Sand+Silt+BD+pH, random=~ phylogeny + species) | (Intercept) | -0.1793 | -0.9490 | 0.6068 | 1700 | 0.6447 | R^2^m = 0.063 |
|  | AI | 0.0994 | -0.2061 | 0.4664 | 1700 | 0.5541 | R^2^p = 0.573 |
|  | MAT | 0.1312 | -0.0365 | 0.2800 | 947.3 | 0.1024 | R^2^s = 0.079 |
|  | **TS** | **-0.3240** | **-0.5045** | **-0.1456** | **1700** | **0.0012** | R^2^c = 0.715 |
|  | MAP | -0.0177 | -0.2268 | 0.1991 | 1700 | 0.8624 |  |
|  | PS | -0.0245 | -0.1450 | 0.1020 | 1700 | 0.7047 |  |
|  | Solar | 0.0413 | -0.1142 | 0.1967 | 1439.7 | 0.5718 |  |
|  | N | 0.0700 | -0.0657 | 0.2254 | 1700 | 0.3377 |  |
|  | Sand | 0.0653 | -0.1625 | 0.2846 | 1657.8 | 0.5706 |  |
|  | Silt | 0.1090 | -0.0954 | 0.3516 | 1538.7 | 0.3141 |  |
|  | BD | 0.0096 | -0.1160 | 0.1238 | 1477.1 | 0.8941 |  |
|  | pH | 0.1513 | -0.0488 | 0.3618 | 1535.5 | 0.1447 |  |

The abbreviations of environmental factors were shown in Table S1. Significant effects of environmental factors on stomatal traits are indicated in bold. Post.mean, posterior mean; Eff.samp, the effective sample size; pMCMC, p-value from Monte Carlo sampling by Markov Chain. Lower 95% CI, the lower limit of the 95% confidence interval for slope. Upper 95% CI, the upper limit of the 95% confidence interval for slope. R^2^c, percentage of variance explained by all the model (fixed + random); R^2^m, percentage of variance explained by fixed factors; R^2^p, percentage of variance explained by phylogeny; R^2^s, percentage of variance explained by species.

Bold values indicate that the 95% credible intervals exclude zero and, therefore, are statistically significant.

**Table S21 Results from Bayesian phylogenetic linear mixed model of the stomatal ratio (SR), with fixed factors (i.e., environmental factors) and random factors (i.e., phylogeny + species) taken into account.**

| Bayesian model | Variable | Post.mean | Lower 95% CI | Upper 95% CI | Eff.samp | pMCMC | R^2^ of the model |
| --- | --- | --- | --- | --- | --- | --- | --- |
| MCMCglmm.updateable(SR~AI+MAT+TS+MAP+PS+solar+N+Sand+Silt+BD+pH, random=~ phylogeny + species) | (Intercept) | 0.0629 | -0.6408 | 0.7590 | 1547 | 0.8494 | R^2^m = 0.029 |
|  | AI | 0.0432 | -0.2310 | 0.3247 | 1700 | 0.7753 | R^2^p = 0.638 |
|  | MAT | 0.0149 | -0.1167 | 0.1551 | 1700 | 0.8341 | R^2^s = 0.014 |
|  | **TS** | **0.1999** | **0.0499** | **0.3485** | **1700** | **0.0129** | R^2^c = 0.681 |
|  | MAP | -0.0037 | -0.2044 | 0.1846 | 1700 | 0.9565 |  |
|  | **PS** | **0.1324** | **0.0201** | **0.2482** | **1700** | **0.0271** |  |
|  | Solar | -0.0480 | -0.1781 | 0.0727 | 1548 | 0.4576 |  |
|  | N | 0.0709 | -0.0483 | 0.1970 | 1353 | 0.2753 |  |
|  | Sand | -0.1363 | -0.3541 | 0.0551 | 1700 | 0.2047 |  |
|  | Silt | -0.1039 | -0.2928 | 0.0999 | 1700 | 0.3141 |  |
|  | BD | 0.0702 | -0.0364 | 0.1726 | 1700 | 0.1765 |  |
|  | pH | 0.0363 | -0.1589 | 0.2065 | 1529 | 0.6694 |  |

The abbreviations of environmental factors were shown in Table S1. Significant effects of environmental factors on stomatal traits are indicated in bold. Post.mean, posterior mean; Eff.samp, the effective sample size; pMCMC, p-value from Monte Carlo sampling by Markov Chain. Lower 95% CI, the lower limit of the 95% confidence interval for slope. Upper 95% CI, the upper limit of the 95% confidence interval for slope. R^2^c, percentage of variance explained by all the model (fixed + random); R^2^m, percentage of variance explained by fixed factors; R^2^p, percentage of variance explained by phylogeny; R^2^s, percentage of variance explained by species.

Bold values indicate that the 95% credible intervals exclude zero and, therefore, are statistically significant.

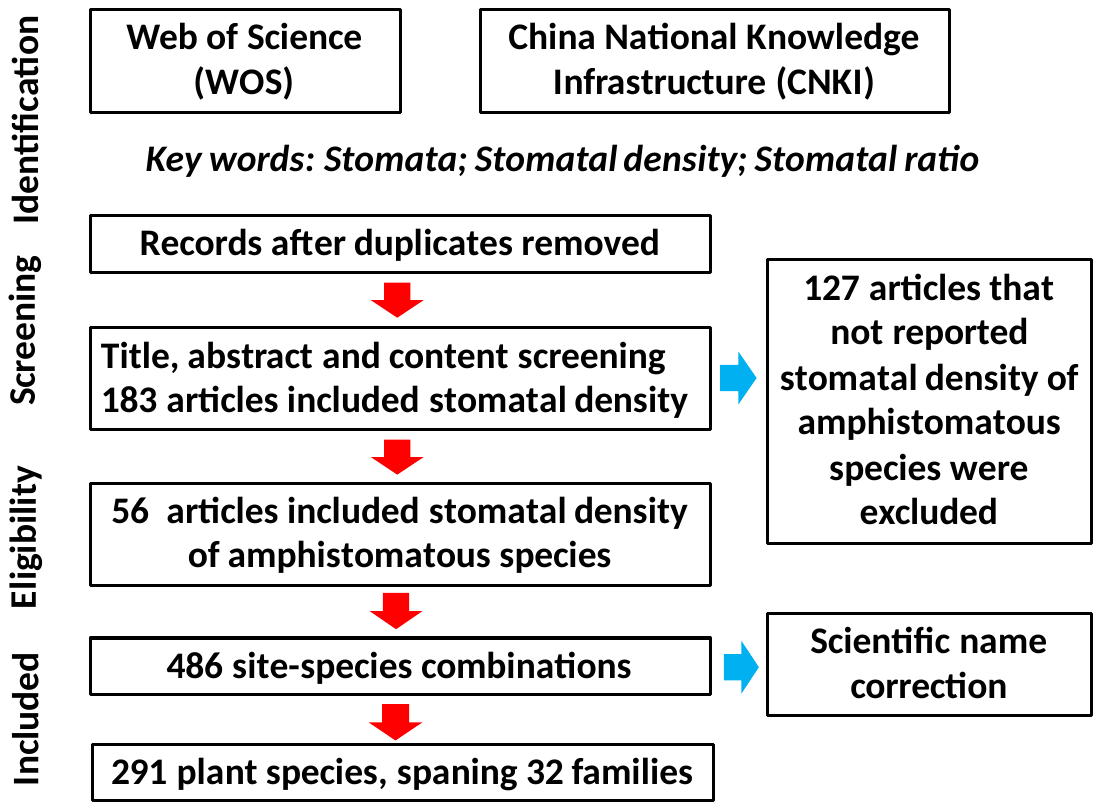

**Fig. S1 The preferred reporting items for systematic reviews and meta-analyses (PRISMA) flowchart of the publication selection procedure.**
